# Regulation of CSF and brain tissue sodium levels by the blood-CSF and blood-brain barriers during migraine

**DOI:** 10.1101/572727

**Authors:** Hamed Ghaffari, Samuel C. Grant, Linda R. Petzold, Michael G. Harrington

**Affiliations:** Department of Mechanical Engineering, University of California Santa Barbara, Santa Barbara, CA, USA; Department of Chemical and Biomedical Engineering, FAMU-FSU College of Engineering, Tallahassee, FL, USA; Molecular Neurology Program, Huntington Medical Research Institutes, Pasadena, CA, USA

**Keywords:** Migraine, Sodium, Blood-brain barrier, Blood-CSF barrier, Mathematical model, Global sensitivity analysis, Sobol’s method, Na^+^, K^+^-ATPase

## Abstract

**Background:** Cerebrospinal fluid (CSF) and brain tissue sodium levels increase during migraine. However, little is known regarding the underlying mechanisms of sodium homeostasis disturbance in the brain during the onset and propagation of migraine. Exploring the cause of sodium dysregulation in the brain is important, since correction of the altered sodium homeostasis could potentially treat migraine. Under the hypothesis that disturbances in sodium transport mechanisms at the blood-CSF barrier (BCSFB) and/or the blood-brain barrier (BBB) are the underlying cause of the elevated CSF and brain tissue sodium levels during migraines, we developed a mechanistic, differential equation model of a rat’s brain to compare the significance of the BCSFB and the BBB in controlling CSF and brain tissue sodium levels. The model includes the ventricular system, subarachnoid space, brain tissue and blood. Sodium transport from blood to CSF across the BCSFB, and from blood to brain tissue across the BBB were modeled by influx permeability coefficients *P*_*cp*_ and *P*_*bc*_, respectively, while sodium movement from CSF into blood across the BCSFB, and from brain tissue to blood across the BBB were modeled by efflux permeability coefficients 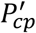 and 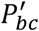, respectively. We then performed a global sensitivity analysis to investigate the sensitivity of the ventricular CSF, subarachnoid CSF and brain tissue sodium concentrations to pathophysiological variations in 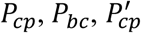 and 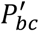. Our results show that the ventricular CSF sodium concentration is highly influenced by perturbations of *P*_*cp*_, and to a much lesser extent by perturbations of 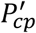. Brain tissue and subarachnoid CSF sodium concentrations are more sensitive to pathophysiological variations of *P*_*bc*_ and 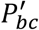 than variations of *P*_*cp*_ and 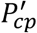 within 30 minutes of the onset of the perturbations. However, *P*_*cp*_ is the most sensitive model parameter, followed by *P*_*bc*_ and 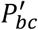, in controlling brain tissue and subarachnoid CSF sodium levels within 2 hours of the perturbation onset.

## 1. Introduction

Migraine is ranked among the top five causes of disability in the world [1]. Although the exact underlying causes of migraine are not known, common triggers of migraine include dehydration, stress, sleep disorders, hunger, etc. Understanding the pathophysiology of migraine is challenging because migraine triggering is different for everyone. Many of the triggers of migraine change the sodium balance in the brain. Previous animal and human studies [2-4] have revealed that migraine sufferers have higher levels of cerebrospinal fluid (CSF) and brain interstitial fluid (ISF) sodium than control groups, while there is no significant difference between blood concentration of sodium in migraineurs and healthy controls. Previous studies have indicated that elevated levels of ISF sodium increase neuronal excitability [5], which subsequently results in migraine [6]. Brain sodium levels ultimately derive from peripheral circulation. Sodium is exchanged between the blood and brain across two major blood-brain interfaces, namely the blood-brain barrier (BBB) and the blood-CSF barrier (BCSFB). The BBB is formed by specialized endothelial cells lining the cerebral microvasculature and controls sodium exchange between the ISF and blood, while the BCSFB is formed by choroid plexus epithelial cells and regulates sodium transport between ventricular CSF and blood. Transfer of sodium across the BBB and the BCSFB predominantly take places via active, hence transcellular mechanisms. However, sodium may be able to cross the BCSFB and the BBB via a paracellular route through tight junctions between epithelial cells at the BCSFB and between endothelial cells at the BBB [7].

It is believed that the BCSFB and BBB are highly responsible for maintaining ion homeostasis in the brain. Thus, a disturbance in sodium transport mechanisms at the BCSFB and/or BBB can alter CSF and brain tissue sodium concentrations. However, the relative contributions of the two interfaces in the regulation of brain sodium homeostasis have yet to be determined. Understanding the roles and significance of the BCSFB and BBB in the regulation of brain tissue and CSF sodium levels is important not only because it helps understand how a migraine starts, but also it potentially offers a new strategy to normalize the elevated brain sodium levels in migraine sufferers and subsequently treat migraines. In this work, we use mechanistic modeling to study the significance of the BCSFB and BBB in controlling brain tissue and CSF sodium levels. We develop a mathematical model consisting of four compartments: the ventricular system, subarachnoid space, brain tissue and blood. Net movement of sodium across the BCSFB and BBB through different active and passive transport mechanisms is modeled by influx and efflux permeability coefficients of the interfaces to sodium. Influx permeability coefficients of the BCSFB and BBB to sodium refer to sodium movement from blood to CSF and brain tissue, respectively, whereas efflux permeability coefficients of the BCSFB and BBB to sodium represent sodium movement from CSF and brain tissue to blood, respectively. We study the dynamics of sodium distribution in the brain following a perturbation in the influx and efflux permeabilities of the BCSFB and BBB to sodium. We then perform a global sensitivity analysis (GSA) to assess the significance of the BCSFB and BBB in controlling sodium concentrations in the brain tissue, ventricular CSF and subarachnoid CSF. Our results reveal that the influx permeability coefficient of the BCSFB to sodium is the most sensitive model parameter in controlling ventricular CSF sodium concentration. Depending on the time elapsed from perturbations of the permeability coefficients, brain tissue and subarachnoid space CSF sodium levels can be significantly controlled by the BCSFB and/or BBB.

The computational model presented in this study can not only shed light on the dynamics of sodium exchange between CSF, brain tissue and blood, but can also provide insight for future experimental studies. In addition, this work can potentially offer a new strategy to normalize the elevated levels of brain sodium in migraine sufferers and potentially treat migraines.

## 2. Methods

### 2.1. Model Development

We modeled a rat’s brain by three concentric spheres representing the ventricular system, brain tissue and subarachnoid space (Fig. 1). Brain tissue was modeled as a single compartment. We assumed that blood vessels are distributed randomly, following a uniform distribution, throughout the brain tissue[8-11].

**Figure 1.**
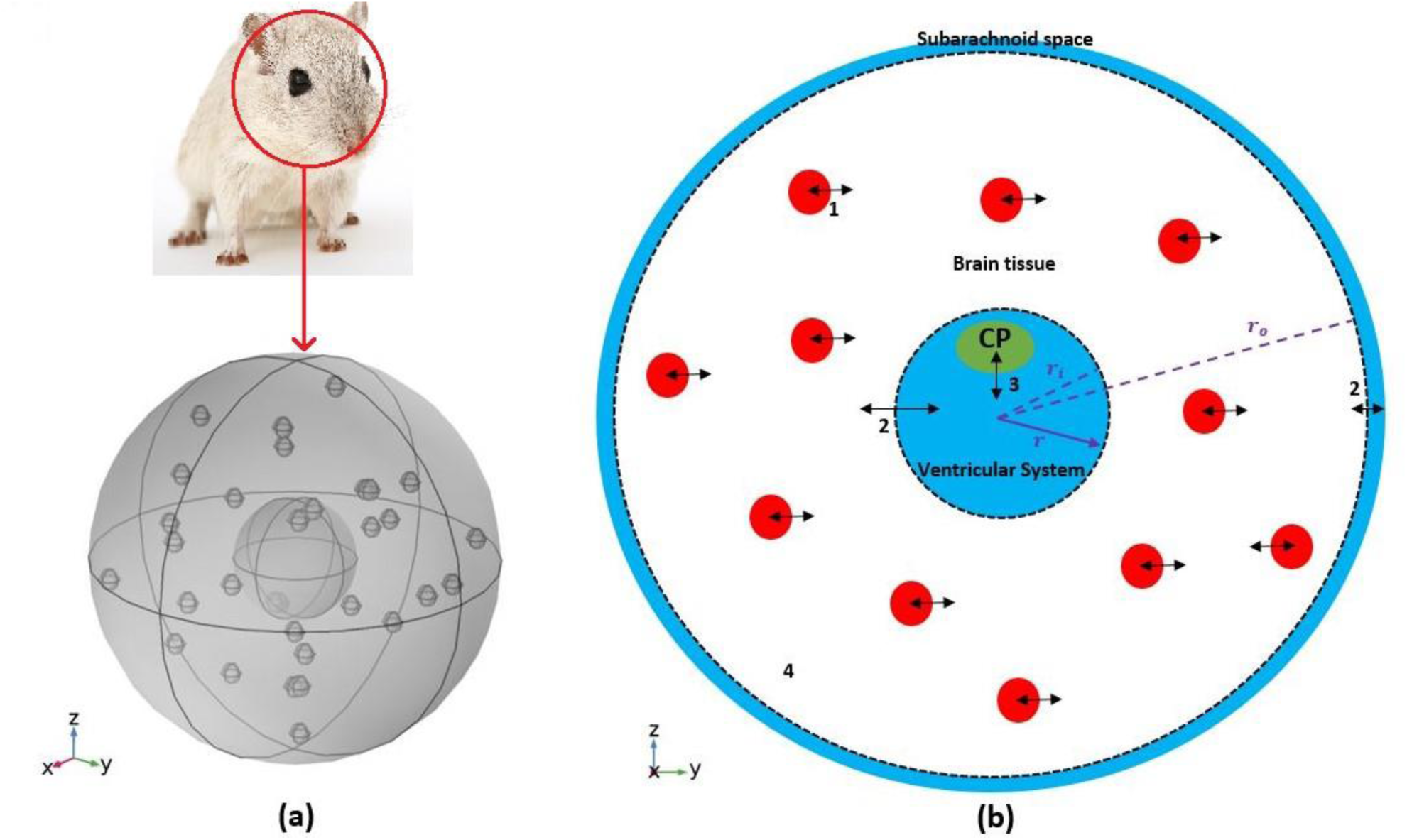
Schematic of the model. (a) A 3D model of a rat’s brain. (b) A 2D view of the cross section of the 3D model. The inner circle, shown in blue, represents the ventricular system, while the outer ring, shown in blue, is subarachnoid space. The white region between two dashed circles is brain tissue. Blood vessels, shown in red circles, are distributed uniformly in the brain tissue. The CP is depicted by a green ellipsoid. Numbers in the figure specify the types and locations of sodium transport: 1. capillary-brain transport; 2. free exchange between CSF and ISF; 3. blood-CSF exchange at the CP; 4. diffusive transport across brain ISF.

The inner sphere, which represents the ventricular system, includes the CP. CSF is secreted by the CP, flows into the ventricular system, and then passes through small openings (foramina) into the subarachnoid space where it is absorbed through blood vessels into the bloodstream. It has also been suggested that a part of subarachnoid CSF moves into the brain along paravascular routes surrounding cerebral arteries, where it mixes with brain ISF and leaves the brain along veins [12, 13]. In the current model, we have ignored CSF flow from subarachnoid space to brain ISF (See Section 4 for further discussion of this subject). Thus, we have assumed that the CSF secretion rate is equal to the CSF absorption rate from the subarachnoid space to the blood. We have also assumed that sodium can be freely exchanged between the brain tissue and the CSF at the interface of brain tissue and the ventricular system, and at the contact surface of the subarachnoid space and brain tissue (dashed circles in Fig. 1b). This is due to the negligible permeability barrier of the contact surfaces. Sodium is also exchanged between blood and brain tissue across the BBB, and can also diffuse in the brain tissue down its concentration gradient.

### 2.2. Formulation of the model

Ventricular and subarachnoid CSF sodium concentrations were modeled by ordinary differential equations (ODEs) represented by Eqs. 1-2, while the variation of sodium level across brain tissue was modeled by a partial differential equation (PDE), represented by Eq. 3.

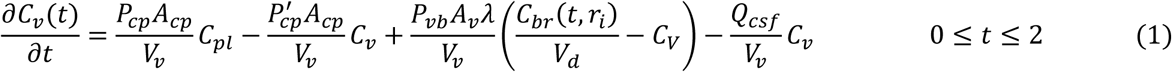

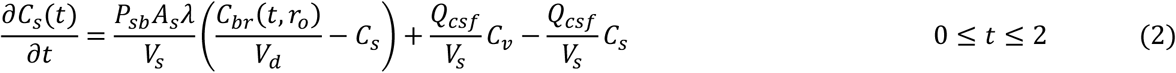

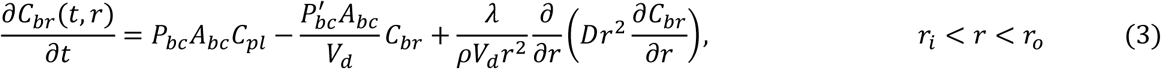

where *C*_*v*_, *C*_*s*_, *C*_*pl*_ and *C*_*br*_ represent ventricular CSF sodium concentration, subarachnoid CSF sodium concentration, blood sodium concentration and sodium level in brain tissue, respectively. *C*_*v*_, *C*_*s*_ and *C*_*pl*_ are expressed in mole ml^−1^, while *C*_*br*_ is defined as moles of sodium per gram of brain (mole g^−1^). *C*_*br*_ includes sodium content in brain ISF and in brain cells. The ISF sodium concentration (mole ml^−1^) was estimated from the brain tissue sodium level (mole g^−1^) by [8]

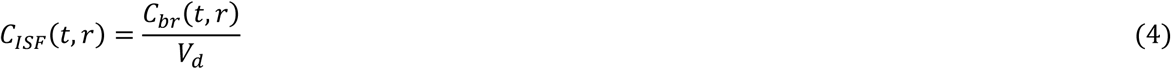

where *V*_*d*_ is sodium distribution factor. The model’s parameters are defined in Table 1. The parameters *r*_*i*_ and *r*_*o*_, which specify the boundaries of brain tissue (Fig. 1b), were obtained via the relationships

**Table 1.** Physiological values of the model’s parameters for an adult rat

| Parameters | Description | Value | Reference |
| --- | --- | --- | --- |
| $P_{cp}$ | BCSFB influx permeability coefficient to sodium (from blood to CSF) | $3.8 \times 10^{-5} \text{ (cm s}^{-1}\text{)}$ | [8] |
| $A_{cp}$ | Surface area of choroid plexus | $1 \text{ (cm}^2\text{)}$ | [8] |
| $P'_{cp}$ | BCSFB efflux permeability coefficient to sodium (from CSF to blood) | Unknown | |
| $V_s$ | Subarachnoid space volume | $0.2 \text{ (cm}^3\text{)}$ | [8, 18] |
| $V_v$ | Ventricular system volume | $0.1 \text{ (cm}^3\text{)}$ | [8, 18] |
| $V_b$ | Brain tissue volume | $1.1 \text{ (cm}^3\text{)}$ | [19] |
| $P_{bc}$ | BBB influx permeability coefficient to sodium (from blood to brain tissue) | $1.4 \times 10^{-7} \text{ (cm s}^{-1}\text{)}$ | [8] |
| $A_{bc}$ | Surface area of the BBB | $140 \text{ (cm}^2 \text{ g}^{-1}\text{)}$ | [8] |
| $P'_{bc}$ | BBB efflux permeability coefficient to sodium (from brain tissue to blood) | Unknown | |
| $V_d$ | Sodium distribution factor | $0.34 \text{ (cm}^3 \text{ g}^{-1}\text{)}$ | [8] |
| $D$ | Diffusion coefficient of sodium in the brain ISF | $1.15 \times 10^{-5} \text{ (cm}^2 \text{ s}^{-1}\text{)}$ | [20] |
| $Q_{csf}$ | CSF flow rate | $3.6 \times 10^{-5} \text{ (cm}^3 \text{ s}^{-1}\text{)}$ | [8] |
| $P_{vb}$ | Permeability coefficient of the contact surface of brain tissue and ventricular system to sodium | $10^6 \text{ (cm s}^{-1}\text{)}$ | A large value was used |
| $P_{sb}$ | Permeability coefficient of the contact surface of brain tissue and subarachnoid space to sodium | $10^6 \text{ (cm s}^{-1}\text{)}$ | A large value was used |
| $\lambda$ | ISF/brain volume fraction | $0.2 \text{ (dimensionless)}$ | [8, 21] |
| $\rho$ | Rat brain density | $1 \text{ (g cm}^{-3}\text{)}$ | [8] |

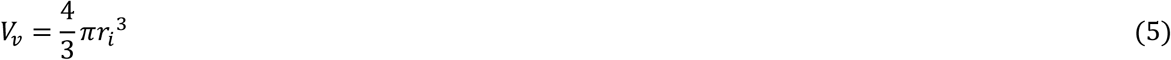

and

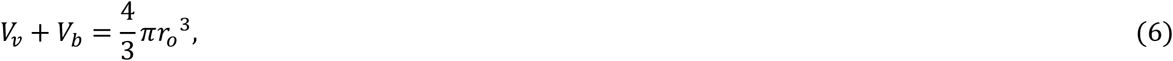

where *V*_*v*_ and *V*_*b*_ represent the ventricular system volume and brain tissue volume, respectively.

The terms on the left-hand side of Eqs. 1 and 2 represent the rate of change of sodium concentration (mole ml^−1^) in the ventricular and subarachnoid CSF, respectively, while the term on the left-hand side of Eq. 3 represents the rate of change of sodium level (mole g^−1^) in the brain tissue. The four terms on the right-hand side of Eq. 1 represent sodium transport from the blood to the ventricular CSF, sodium movement from the ventricular CSF to the blood, sodium exchange between the ventricular CSF and the brain tissue, and sodium loss from the ventricular system due to bulk flow of CSF from the ventricular system to the subarachnoid space, from left to right, respectively. The three terms on the right-hand side of Eq. 2 denote sodium exchange between the subarachnoid CSF and the brain tissue, sodium input to the subarachnoid CSF due to the bulk flow of CSF, and sodium loss from the subarachnoid space due to CSF absorption into the blood, from left to right, respectively. The three terms on the right-hand side of Eq. 3 represent sodium transport from the blood to the brain tissue, sodium movement from the brain tissue to the blood, and diffusive transport of sodium across the brain tissue, from left to right, respectively.

The initial conditions for the ODEs (Eqs. 1-2) are given by [14, 15]

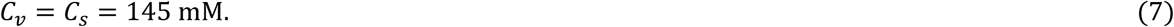

We have also assumed that *C*_*pl*_ is 140 mM at steady state [14].

Rates of exchange of sodium at the boundaries of Eq. 3 are defined by

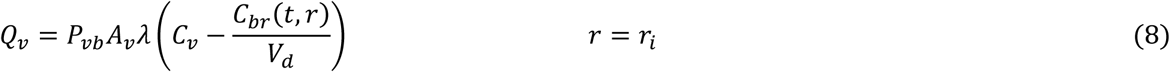

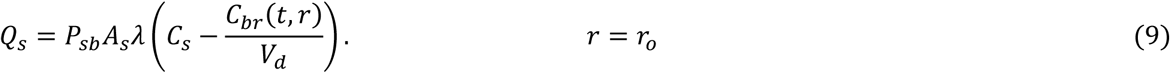

We used large values for *P*_*sb*_ and *P*_*vb*_ due to the negligible permeability barrier of the contact surfaces to sodium. Thus, the ISF sodium concentration is approximately in equilibrium with ventricular and subarachnoid sodium concentrations at the interface of brain tissue and CSF. It is important to note that passive transport of sodium across the boundaries of brain tissue and CSF is regulated by the concentration gradient between the CSF and brain ISF (Eqs. 7-8). Brain ISF sodium concentration is estimated from brain tissue sodium level by Eq. 4. *A*_*v*_ and *A*_*s*_ in Eqs. 8 and 9 represent the contact surface area of the brain tissue and the ventricular system, and the contact surface area of the brain tissue and the subarachnoid space, respectively. The contact surfaces were modeled as concentric spheres with the radiuses of *r*_*i*_ and *r*_*o*_ (Fig. 1). *A*_*v*_ and *A*_*s*_ were obtained by

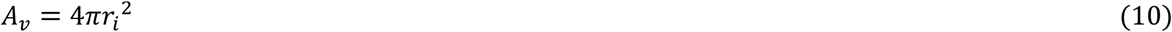

and

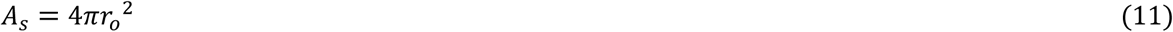

where *r*_*i*_ and *r*_*o*_ were calculated from Eqs. 5 and 6 using the physiological values of *V*_*v*_ and *V*_*b*_ (Table 1). In this model, *A*_*v*_ and *A*_*s*_ were obtained to be 1 and 5.5 cm^2^, respectively, consistent with experimental estimates of the contact surfaces areas [16, 17].

Equation 3 was coupled with Eq. 1 and Eq. 2 at the boundaries of brain tissue (*r* = *r*_*i*_ and *r* = *r*_*o*_), using Eq. 8 and Eq. 9, respectively.

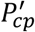 and 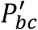 were calculated assuming that the CSF sodium level is in equilibrium with the brain tissue sodium concentration at t=0 (steady state):

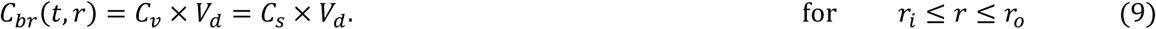

This assumption implies that there is no sodium exchange between the CSF and the brain tissue at the two contact surfaces of brain tissue and CSF at t=0 [22, 23]. The obtained values for 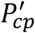 and 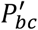 were 6.9 × 10^−7^ cm s^−1^ and 1.35 × 10^−7^ cm s^−1^, respectively. In order to assess the validity of the obtained value for 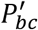, we calculated the rate constant for total sodium efflux from the brain tissue to the blood, defined by 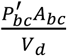 [24]. The average value of 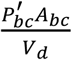 was 5.5 × 10^−5^ s^−1^ in this work, which is consistent with the value of 1 × 10^−4^ s^−1^ reported by Cserr et al [24].

In Section 3, we perform a local sensitivity analysis to investigate how perturbations in 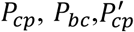 or 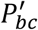 affect brain and CSF sodium concentrations. We also perform a global sensitivity analysis (GSA) to further analyze the significance of variations in the permeability coefficients in controlling the levels of sodium in the CSF and brain tissue. To solve the system of differential equations described by Eqs 1-3, we discretize Eq. 3 with respect to the variable *r* using the central difference approximation, and we approximate the time derivatives via backward differences. The main advantage of this fully implicit scheme, a.k.a. backward time central space, is that it is unconditionally stable.

### 2.3. Global sensitivity analysis

Global sensitivity analysis (GSA) is a numerical method designed to analyze the impacts of uncertain parameters on a model’s output. Compared to local sensitivity analysis, which assesses the changes of model response by making small perturbations to each parameter while keeping the remaining parameters unchanged, GSA analyzes the variations in the model output when all model parameters can vary simultaneously over specified ranges. In other words, GSA investigates how the uncertainty of the model’s output is apportioned to variations in multiple model inputs. This feature makes GSA useful for understanding the contributions of uncertain model parameters to the variations of the model output. In this work, we use GSA to compare the importance of 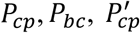 and 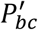 in controlling brain tissue and CSF sodium concentrations, while taking into account the inter-subject variability in all of the model’s parameters. We use a MATLAB toolbox for GSA, called SAFE [25]. We perform Sobol’s sensitivity analysis, which quantitively ranks the relative importance of the parameters by decomposing the model’s output variance into the contributions associated with each model’s input. Sobol’s method, which has been widely applied to complex systems biology and pharmacology models [26-31], calculates the first-order and total-effect sensitivity indices for each model parameter. The first-order indices (*S*_*i*_) measure the individual contributions of each input to the variance of the model output, while the total-effect indices (*S*_*Ti*_) represent the total contribution of the input, including its first-order effect and all higher-order interactions. The total-effect sensitivity indices can be used to identify unimportant model parameters. Non-influential parameters can be fixed at any value within their range of variability without significantly affecting the model response. In Sobol’s sensitivity analysis technique, the model parameters that have total-effect sensitivity indices below 0.01 are often considered non-influential [32, 33] (see Supplementary Information for further details).

## 3. Results

It is believed that brain sodium homeostasis is highly regulated by the BCSFB and the BBB. Elevated levels of sodium in the CSF and brain tissue of migraine sufferers can be due to variations in the influx and/or efflux permeability coefficients of the BCSFB and/or the BBB to sodium. Heuristically, one may expect that the elevated CSF sodium concentration is due to increased transport of sodium from blood into CSF and/or decreased uptake of sodium from CSF into blood. Figure 2 shows the variations in brain tissue, ventricular and subarachnoid CSF sodium concentrations within 2 hours after either a 20% increase in the influx permeability coefficient of the BCSFB to sodium (*P*_*cp*_), or a 20% decrease in the efflux permeability coefficient of the BCSFB to sodium 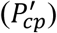.

**Figure 2.**
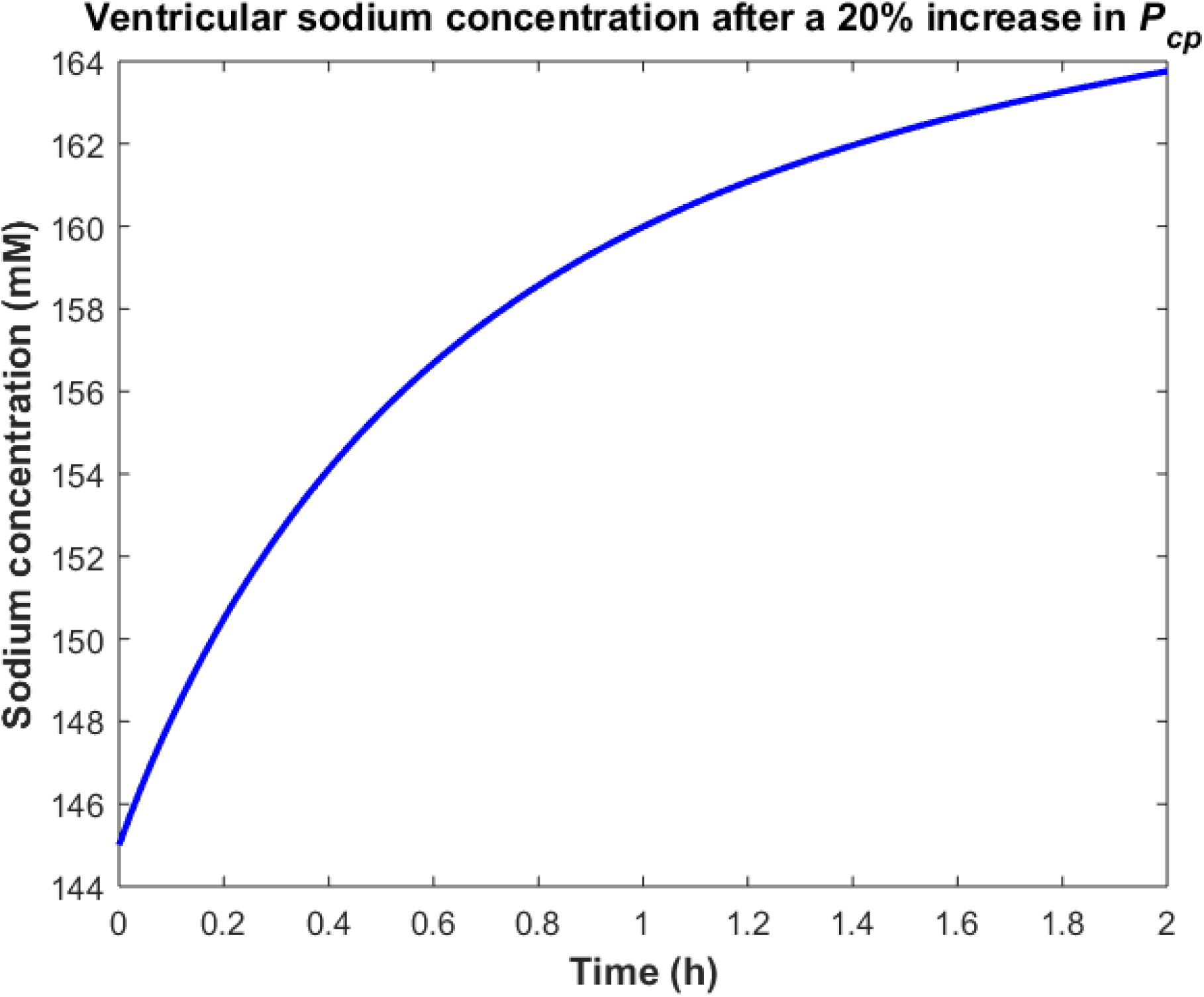
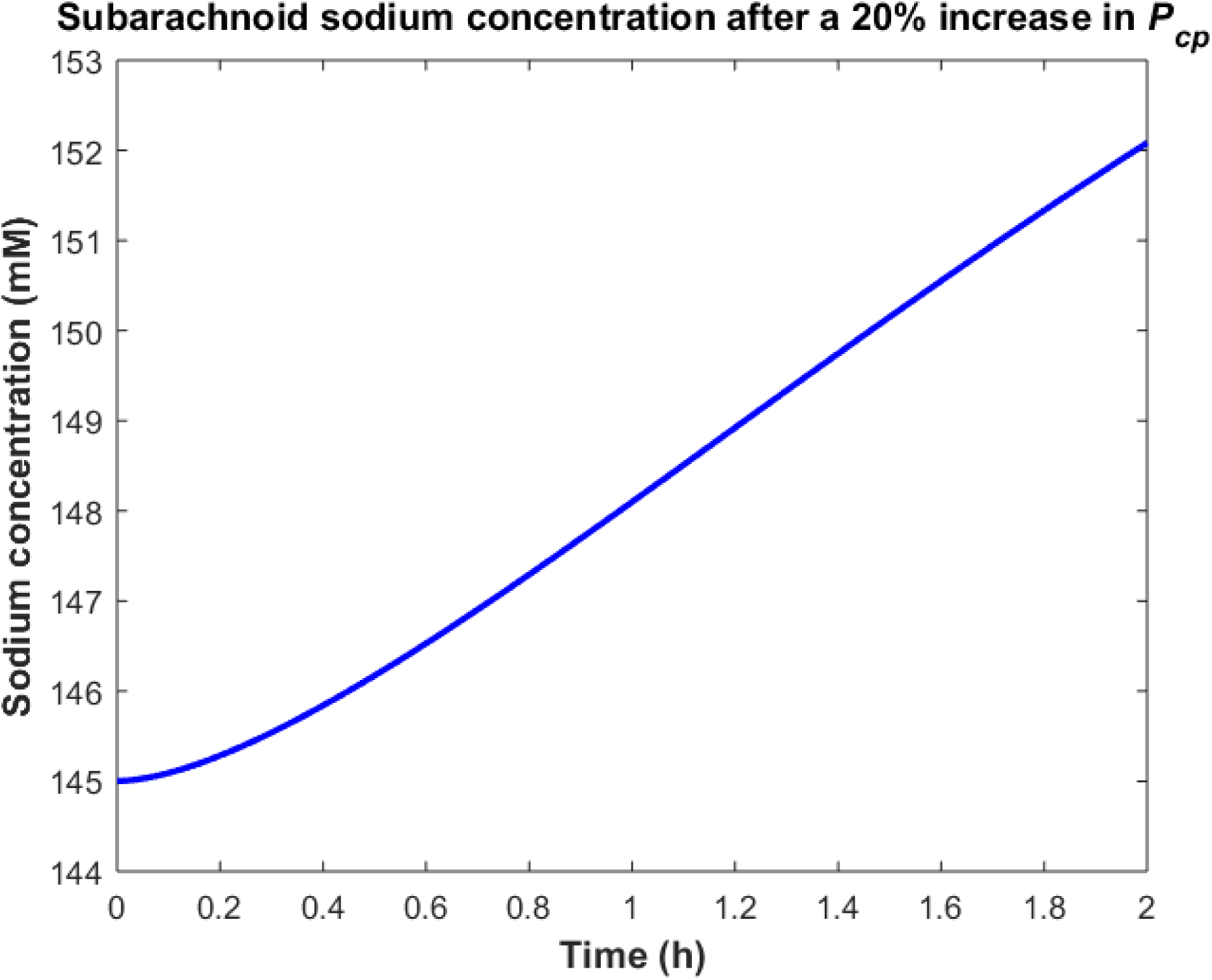
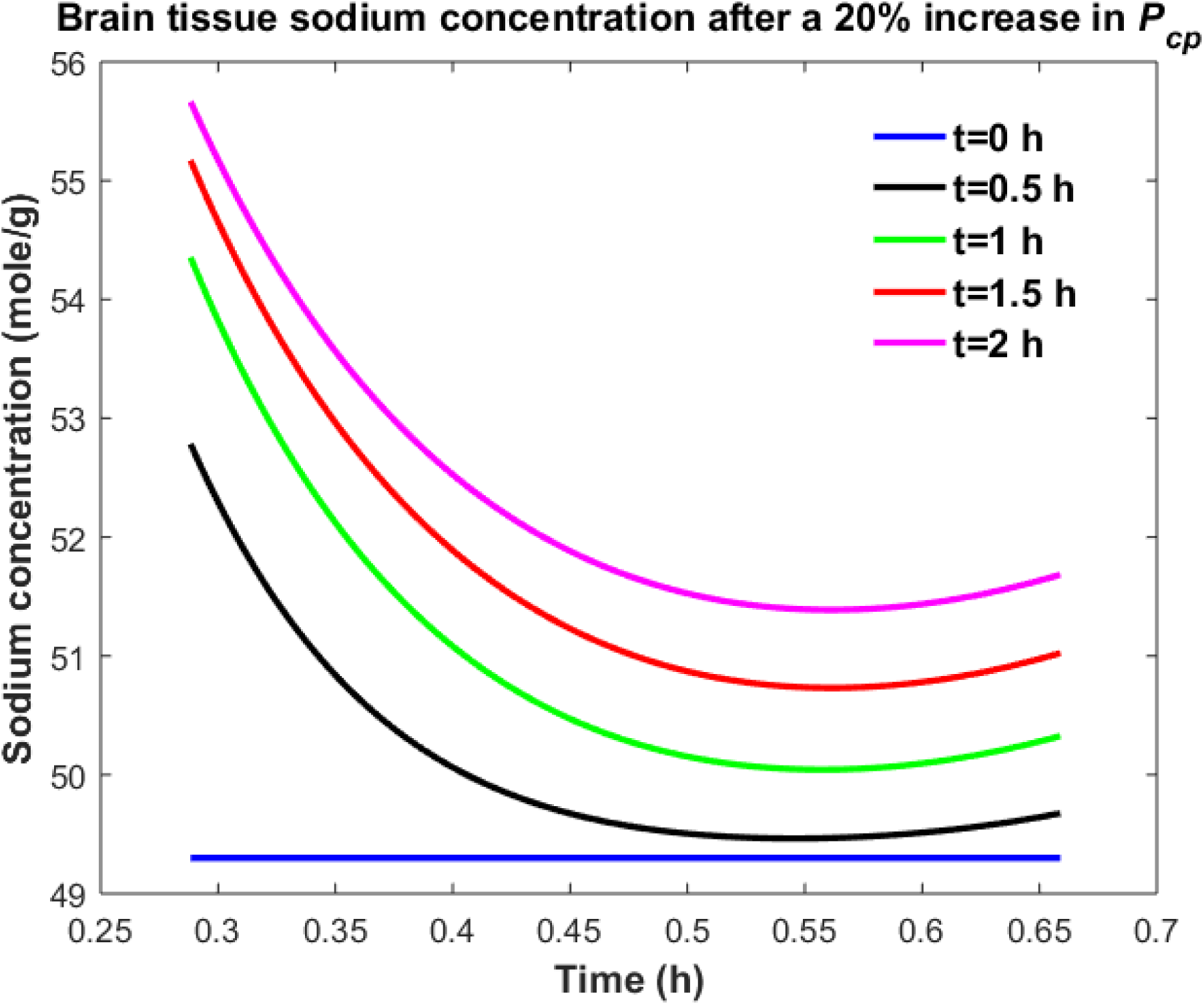
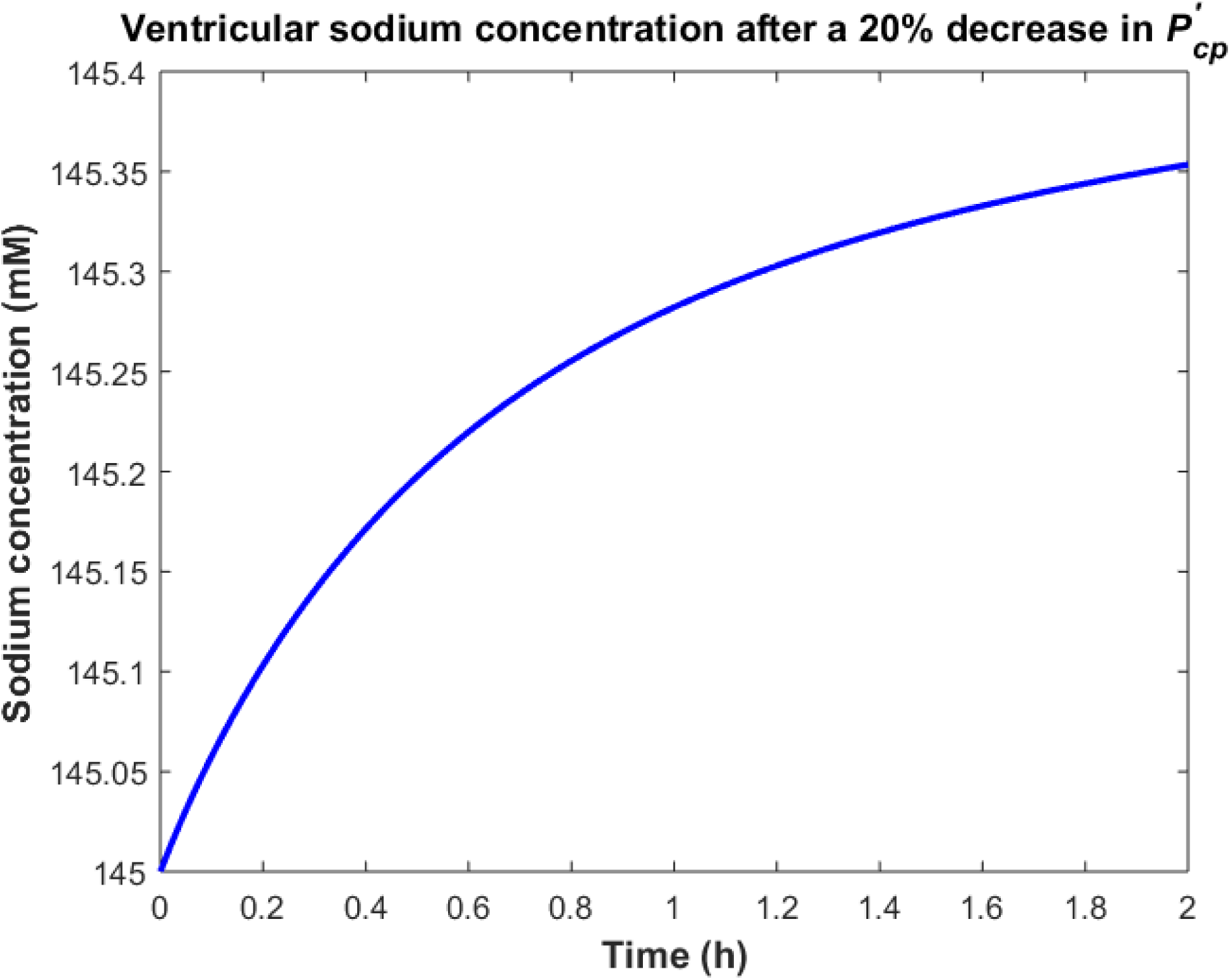
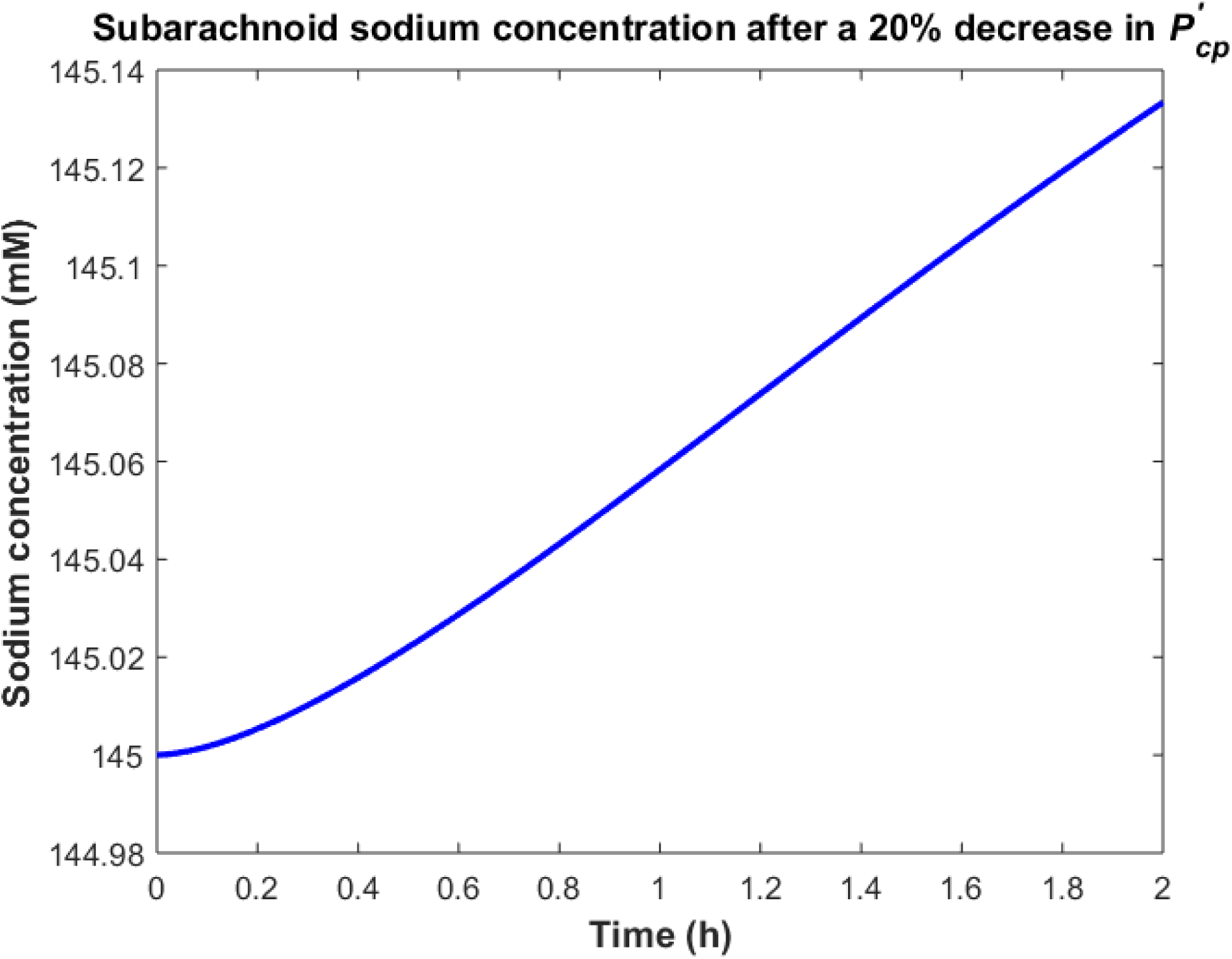
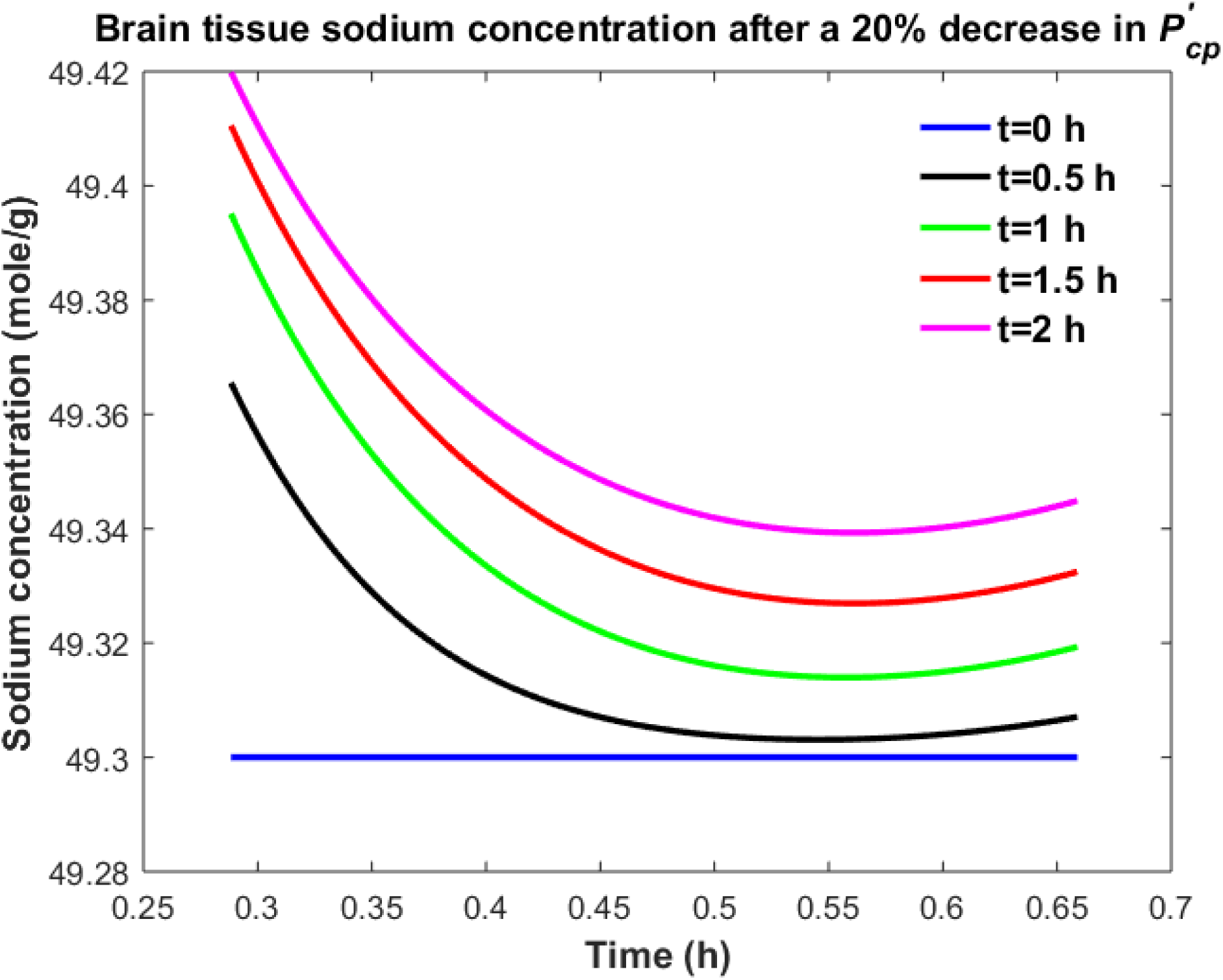
Variations of (a) *C*_*v*_ after increasing *P*_*cp*_ by 20%, (b) *C*_*s*_ after increasing *P*_*cp*_ by 20%, (c) *C*_*br*_ after increasing *P*_*cp*_ by 20%, (d) *C*_*v*_ after decreasing 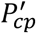 by 20%, (e) *C*_*s*_ after decreasing 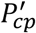 by 20% (f) *C*_*br*_ after decreasing 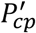 by 20%.

Ventricular CSF sodium concentration increases after a 20% rise in *P*_*cp*_ or a 20% decrease in 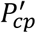 (Figs. 2a and 2d). We assumed that sodium exchange between blood and CSF does not change plasma concentration of sodium significantly, due to the large volume of blood compared to CSF. Thus, *C*_*pl*_ remains unchanged after changing the influx or efflux permeability coefficients of CP to sodium. Figures 2c and 2f show that the elevated levels of sodium in the ventricular CSF lead to diffusion of sodium from CSF to brain tissue and distribution of sodium into the brain tissue over time [8, 34]. Sodium moves by bulk flow of CSF from the ventricular system to the subarachnoid space, where it can be exchanged between CSF and brain tissue. Subarachnoid CSF sodium concentration increases after increasing *P*_*cp*_ or decreasing 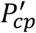 by 20% (Figs. 2b and 2e). Our results indicate that ventricular CSF, subarachnoid CSF and brain tissue sodium levels are more sensitive to variations of *P*_*cp*_ than of 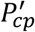. This is because steady state loss of ventricular CSF sodium is largely due to bulk flow of CSF from the ventricular system toward the subarachnoid space rather than to sodium uptake by blood across the BCSFB (Eq. 1 and physiological data in Table 1). However, the only source for sodium in the ventricular system is the choroid plexus at steady state. Thus, a 20% decrease in 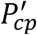 has a less significant impact than a 20% increase in *P*_*cp*_ on CSF sodium content. It should be noted that we assume that there is no sodium exchange between the ventricular CSF and brain tissue at steady state (t=0). Supplementary animations 1 and 2 show the variations of brain ISF, ventricular and subarachnoid sodium concentrations within 2 hours after increasing *P*_*cp*_ or decreasing 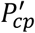 by 20%, respectively.

Similarly, one may expect that the elevated brain tissue sodium levels during migraine are due to increased sodium transport from blood to brain tissue and/or reduced sodium uptake from brain tissue into blood. Figure 3 depicts the changes in ventricular CSF, subarachnoid CSF and brain tissue sodium levels within 2 hours of either increasing the influx permeability coefficient of the BBB to sodium (*P*_*bc*_) by 20%, or decreasing the efflux permeability coefficient of the BCSFB to sodium 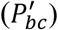 by 20%.

**Figure 3.**
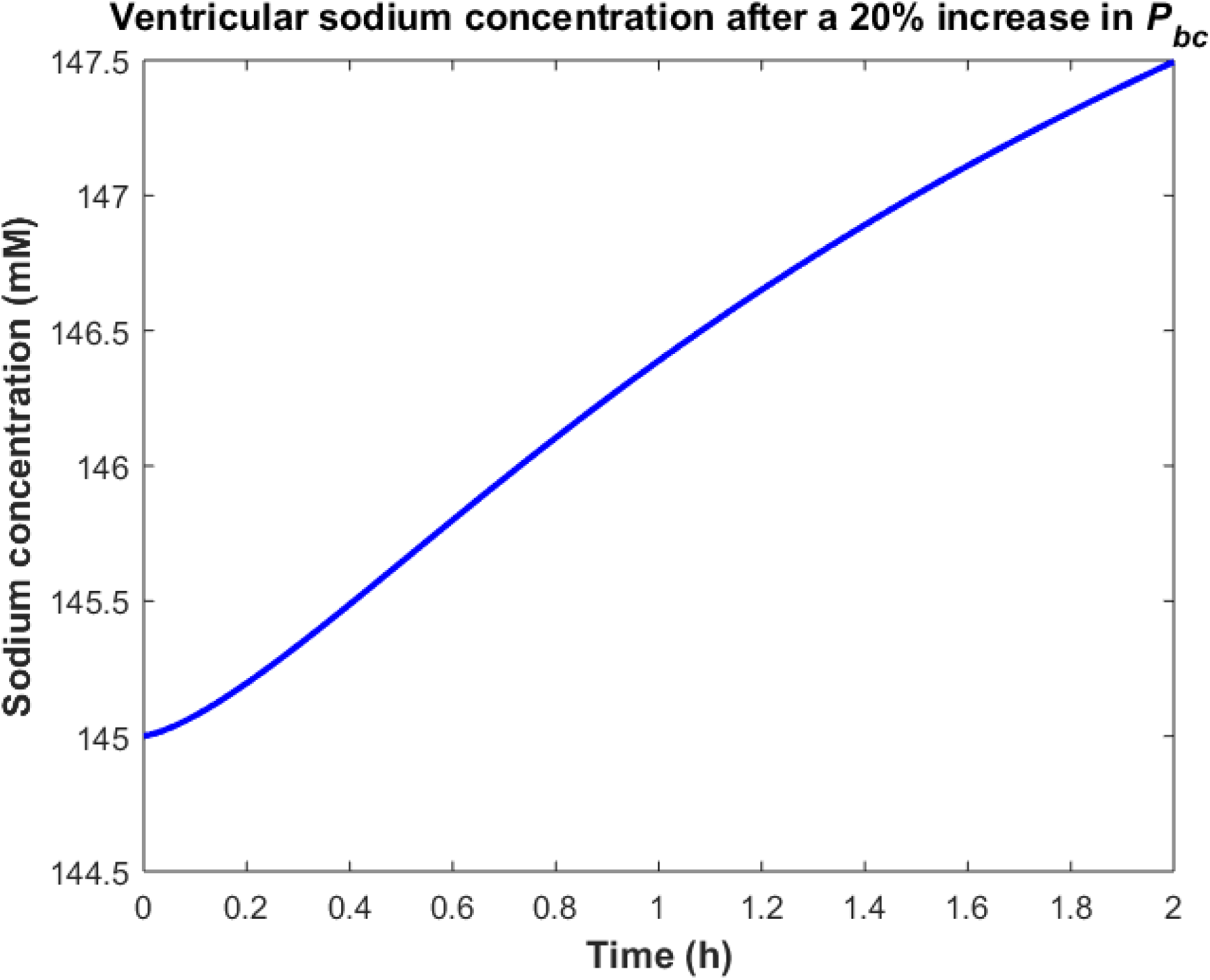
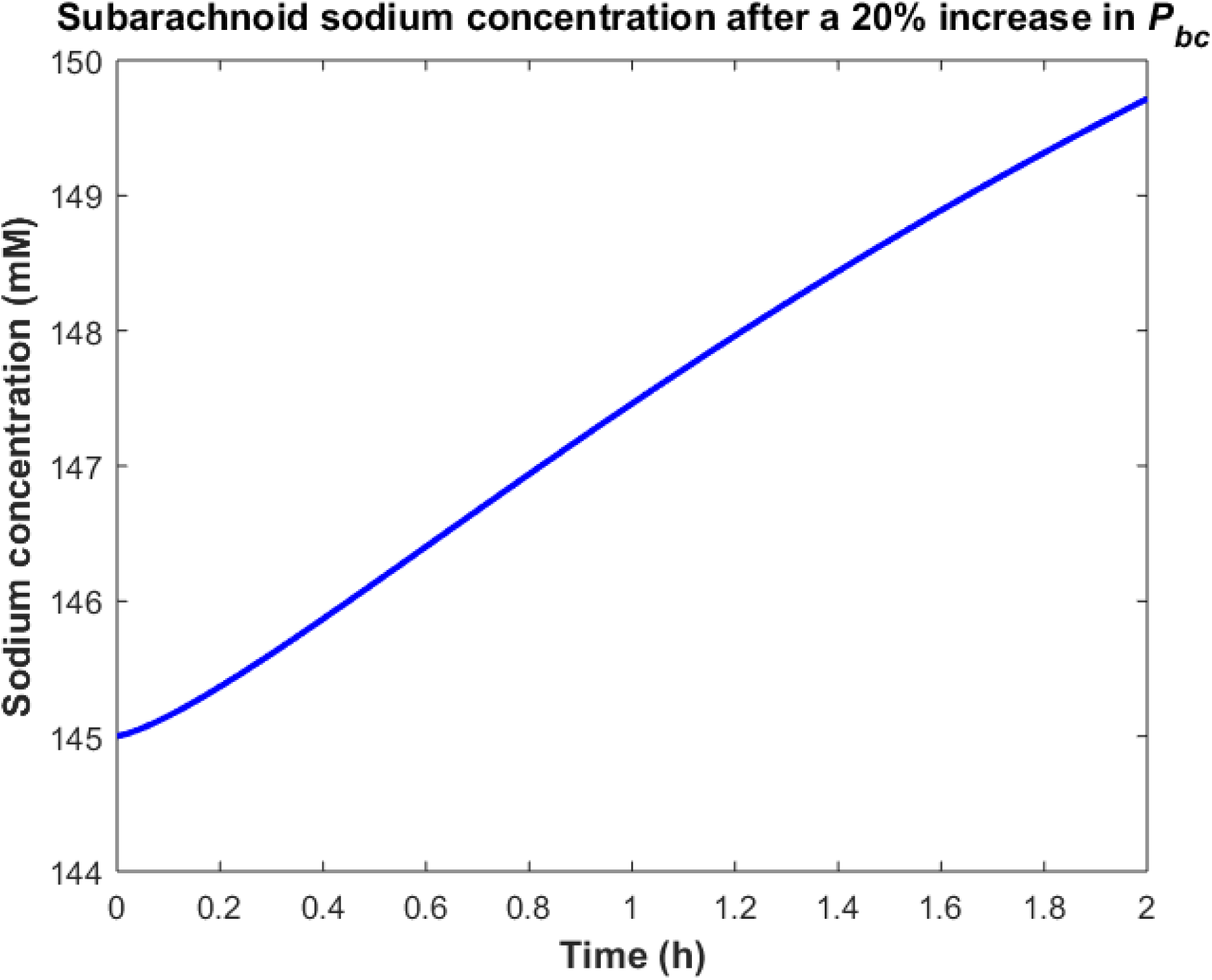
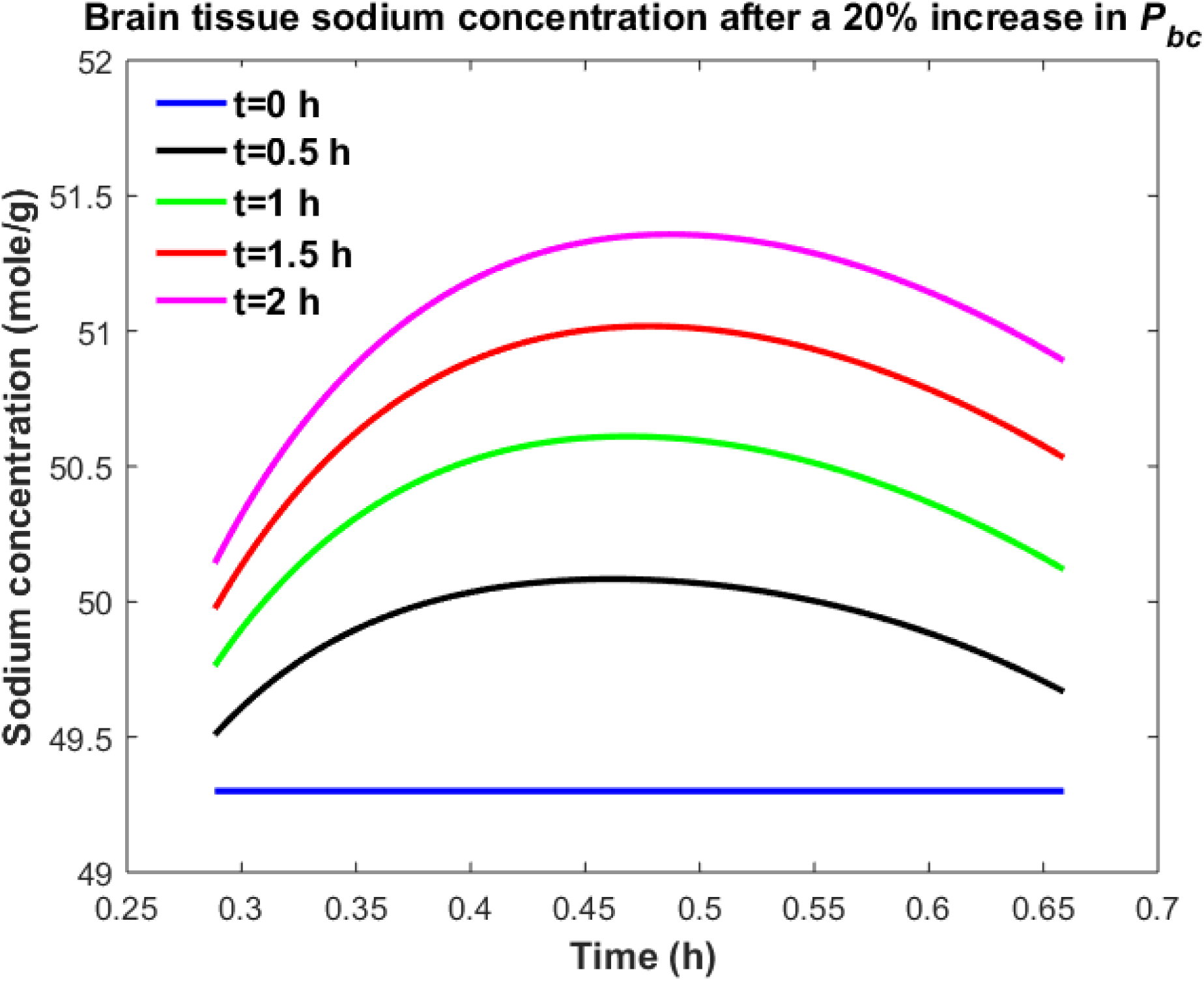
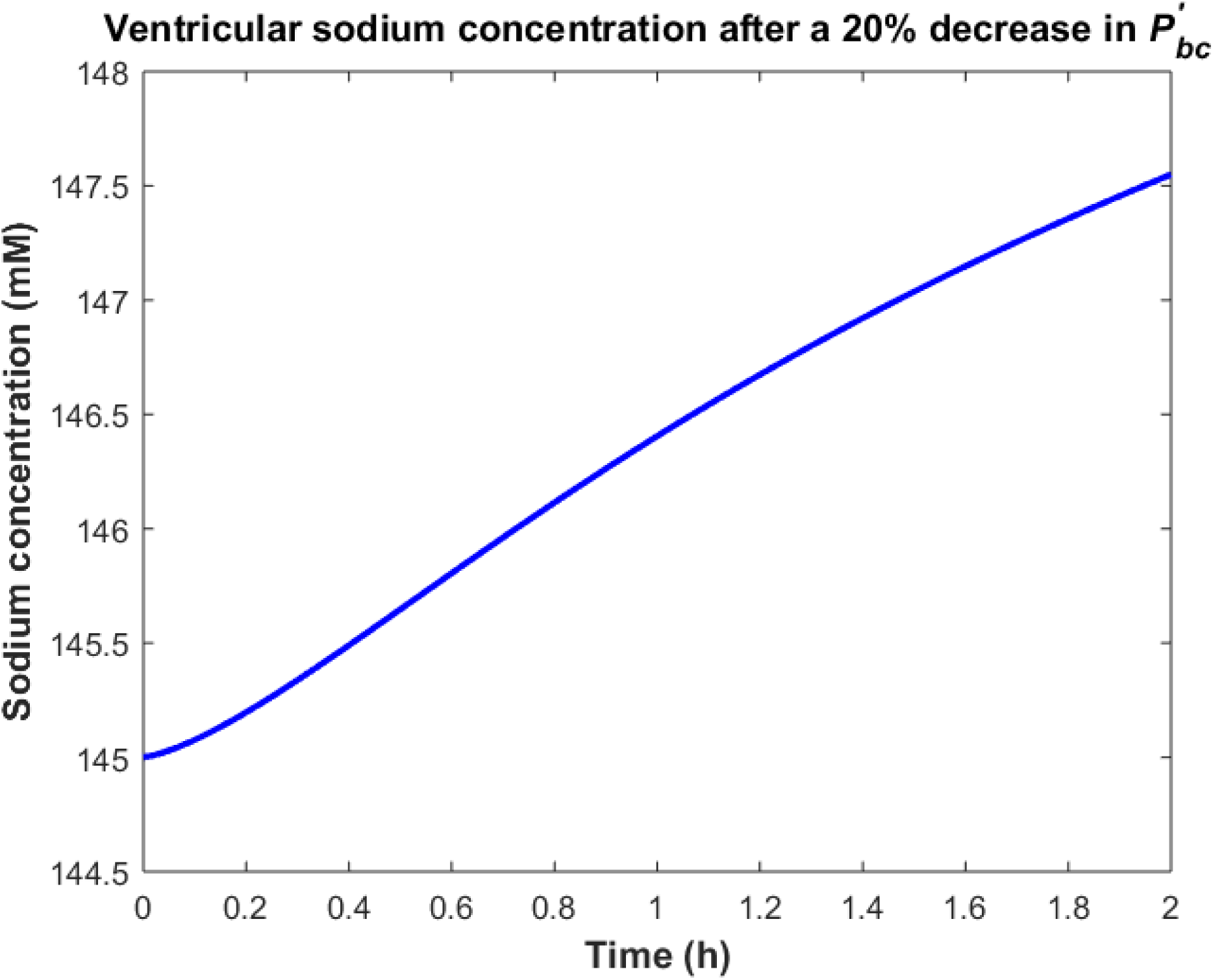
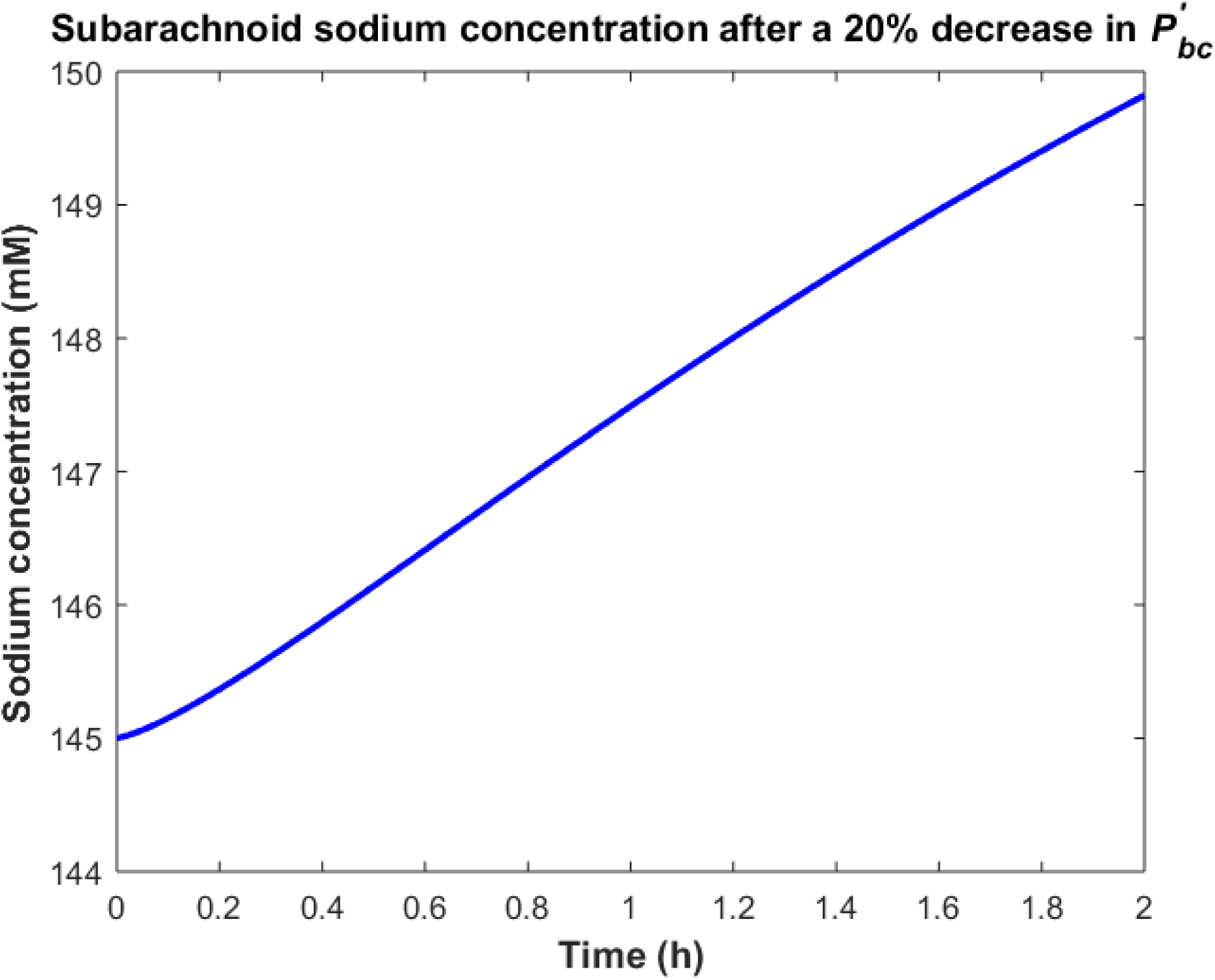
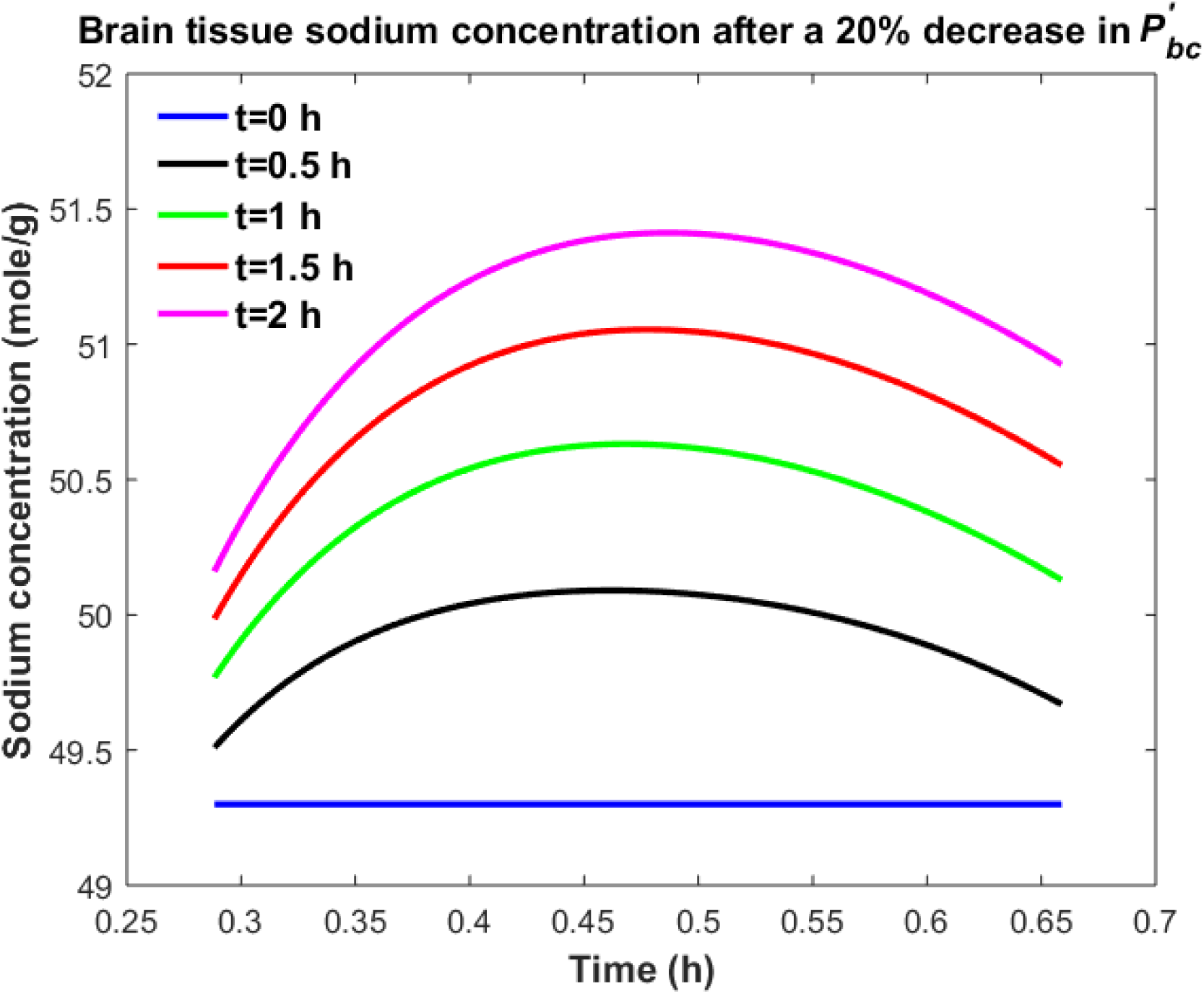
Variations of (a) *C*_*v*_ after increasing *P*_*bc*_ by 20%, (b) *C*_*s*_ after increasing *P*_*bc*_ by 20%, (c) *C*_*br*_ after increasing *P*_*bc*_ by 20%, (d) *C*_*v*_ after decreasing 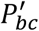 by 20%, (e) *C*_*s*_ after decreasing 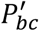 by 20% (f) *C*_*br*_ after decreasing 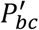 by 20%

A 20% increase in *P*_*bc*_ or a 20% decrease in 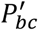 results in an accumulation of sodium in the brain tissue (Figs. 3c and 3f). The elevated levels of sodium in the brain tissue increase sodium transport from brain tissue to the ventricular system and subarachnoid space (Figs. 3a, 3b, 3d and 3e). Our results indicate that brain tissue, ventricular CSF and subarachnoid CSF sodium levels are almost equally sensitive to variations in *P*_*bc*_ and 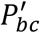. Supplementary animations 3 and 4 show the changes in brain ISF, ventricular and subarachnoid sodium concentrations within 2 hours after increasing *P*_*bc*_ or decreasing 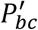 by 20%, respectively.

Figure 4 shows the sodium flux between the brain tissue and CSF at the interface of brain tissue and the ventricular system, and at the contact surface of brain tissue and the subarachnoid space, after perturbation of 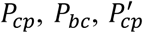 or 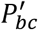 by 20%. Our results indicate that sodium flux from the ventricular system to the brain tissue is larger than sodium flux from the subarachnoid space to the brain tissue.

**Figure 4.**
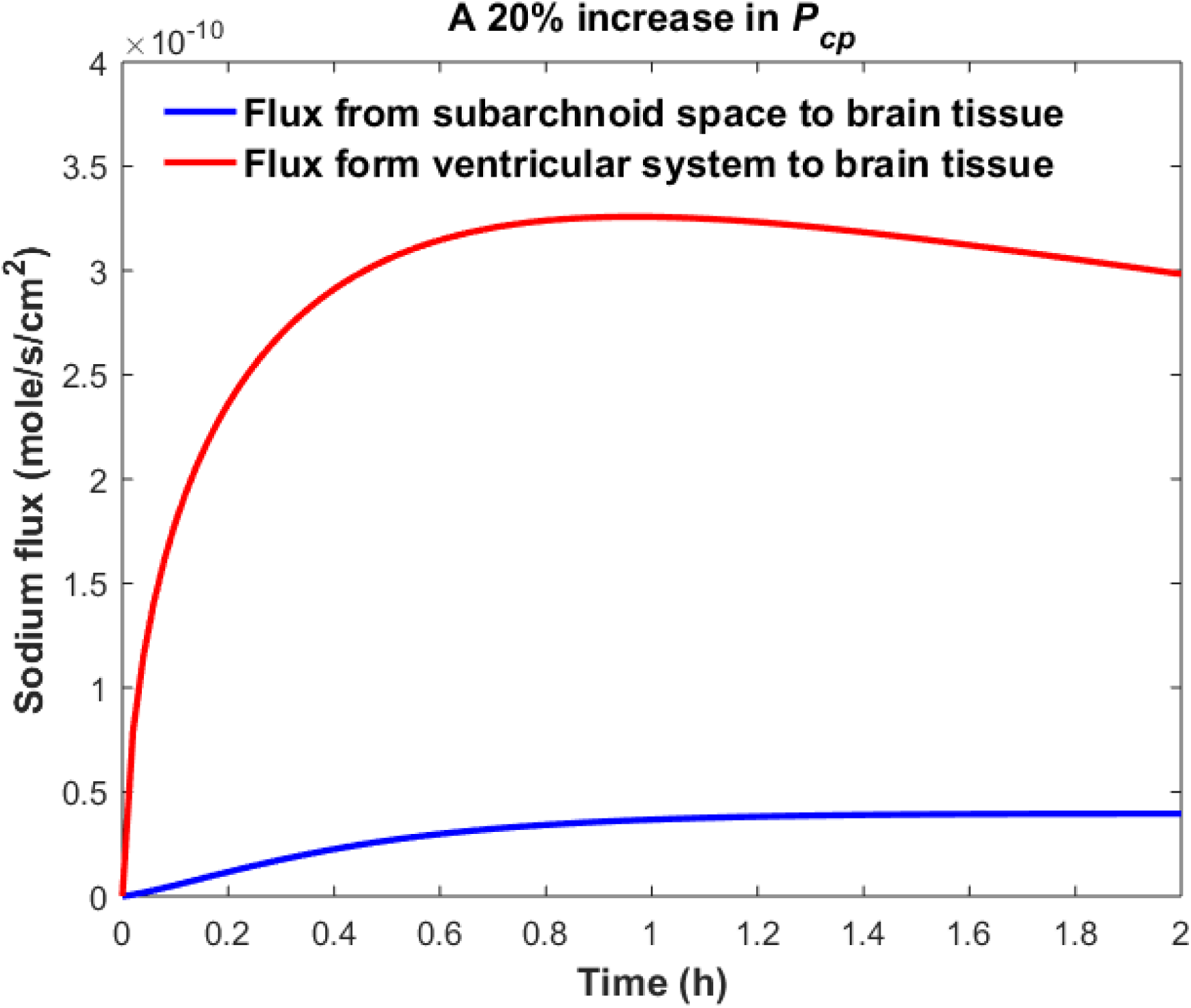
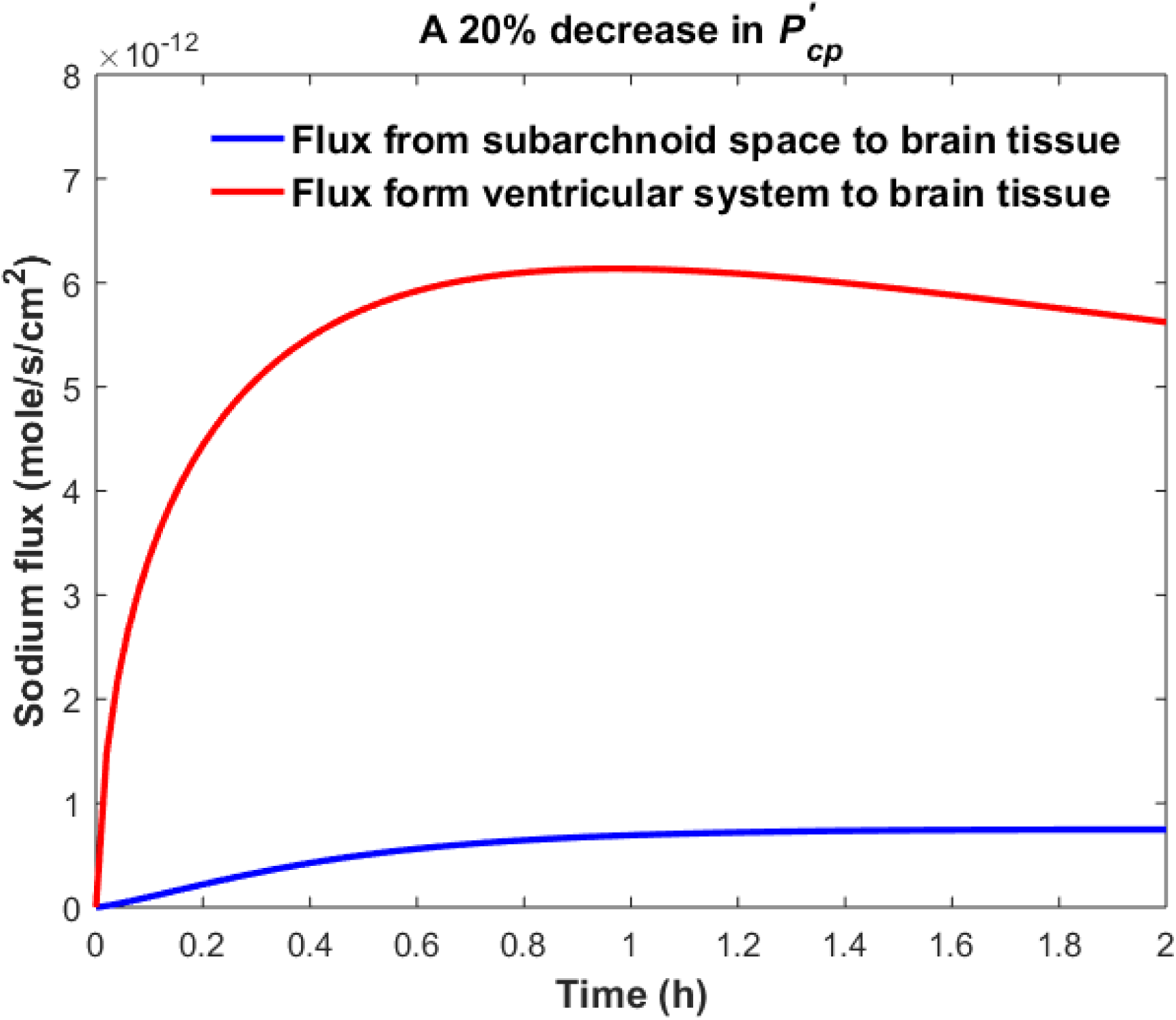
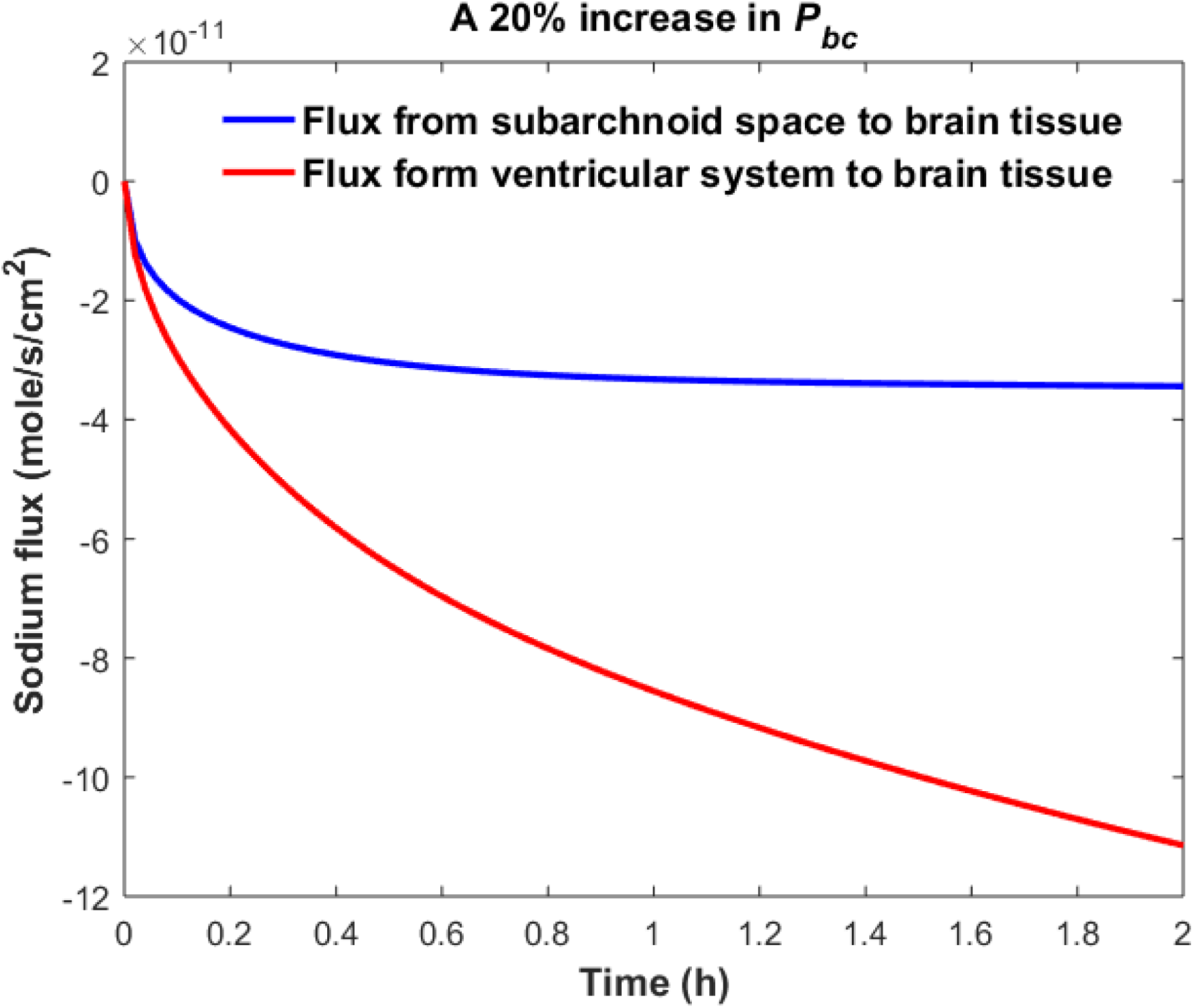
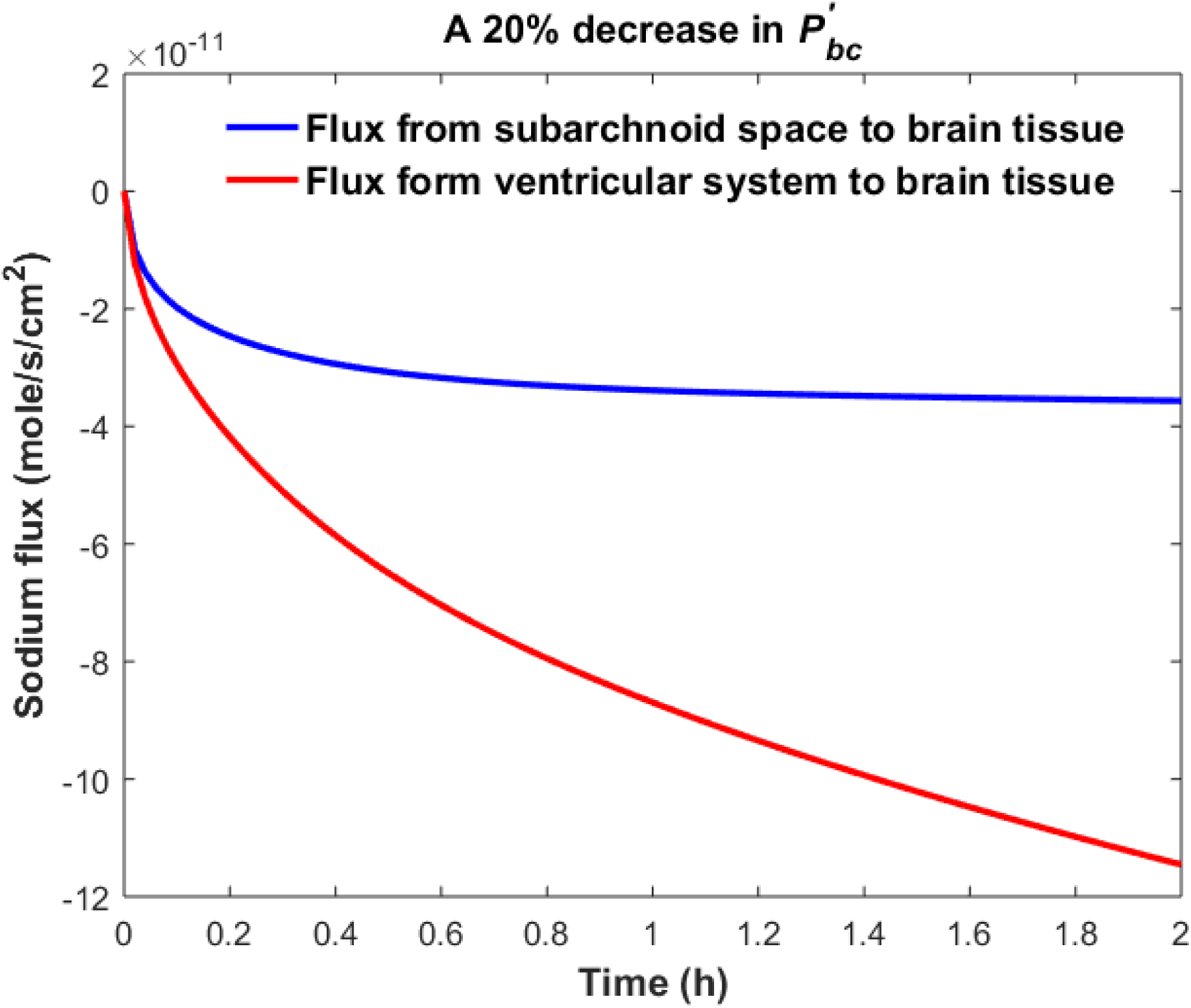
Comparison of sodium flux at the interface of the brain tissue and the ventricular system with sodium flux at the interface of the brain tissue and the subarachnoid space after (a) increasing *P*_*cp*_, (b) Decreasing 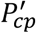, (c) increasing *P* (d) decreasing 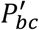 by 20%. The positive sign of the flux indicates that sodium is diffusing from the CSF to the brain tissue, while the negative sign indicates that sodium is diffusing from the brain tissue to the CSF.

Figure 2 and Figure 3 compare the variations in *C*_*v*_, *C*_*s*_, and *C*_*br*_ when a single parameter (i.e. 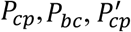 or 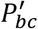) is perturbed and the rest of the parameters remain unchanged. However, in the case of migraines, all influx and efflux permeability coefficients can potentially vary. Additionally, Table 1 shows the average values of the physiological model’s parameters. These values can change across a population of rats of the same type. Thus, we used GSA [25] to consider the effects of variations in all model parameters. In this regard, we assumed that physiological concentration of sodium in CSF and blood can vary within 5% of the *in vitro* values (i.e. *C*_*v*_ = *C*_*s*_ = 145 mM, *C*_*pl*_ = 140 mM), while the remaining independent model parameters (*P*_*cp*_, *A*_*cp*_, *V*_*s*_, *V*_*v*_, *V*_*b*_, *P*_*bc*_, *A*_*bc*_, *V*_*d*_, *D, Q*_*csf*_, *λ, ρ*) can vary within 25% of the *in vitro* values (Table 1). This is due to considering the impacts of intrinsic variations between a population of rats of the same type, and the effects of measurement errors in the estimations of physiological model parameters on our simulations. Following a uniform distribution, we sampled 10^5^ sets of parameters within their ranges of variability. We then calculated the dependent parameters, i.e. 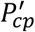 and 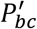 for each set of parameters, assuming that the model is at steady state at t=0. Each of these 10^5^ sets of parameters characterizes one healthy rat with different physiological parameters. We then assumed that 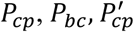, and 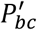 can undergo pathophysiological changes within 50% of their control values due to migraine triggers. We performed a GSA to investigate the significance of pathophysiological variations of 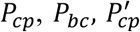, and 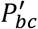 in influencing ventricular sodium concentration during episodic migraines. The model output was defined as the percent change of total ventricular sodium concentration within 2 hours after perturbations of physiological 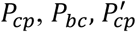, and 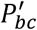:

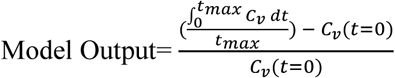

Our results indicate that pathophysiological variation of *P*_*cp*_ is much more important than that of 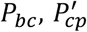, and 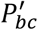 in influencing ventricular CSF sodium concentration (Fig. 5). It is important to note that each permeability coefficient is defined at two states: control and migraine. A given permeability coefficient (e.g. *P*_*cp*_) in the control and migraine states is shown by *P*_*cp*_(*control*) and *P*_*cp*_(*migraine*), respectively. Variations in *P*_*cp*_(*control*) account for intrinsic variations between a population of rats of the same type and/or measurement errors in the estimations of the permeability coefficients. However, a migraine can change physiological permeability coefficients. *P*_*cp*_(*migraine*) represents pathophysiological variations in *P*_*cp*_(*control*) due to migraine triggers. Our results indicate that variations of *P*_*cp*_(*control*) and *P*_*bc*_(*control*) are much less important than that of *P*_*cp*_(*migraine*) and *P*_*bc*_(*migraine*) in influencing the percent change of total ventricular sodium concentration during migraines. This is mainly because the model output was defined as the percent change of total ventricular CSF sodium concentration between the migraine and control states. These results suggest that the ventricular CSF sodium concentration is more sensitive to an alteration in homeostasis of the transporters which mediate sodium influx into CSF across the BCSFB than to a variation in homeostasis of the transporters which regulate sodium uptake from the CSF across the BCSFB. In addition, the BBB plays a much less important role than the BCSFB in regulation of the ventricular CSF sodium concentration. It is important to note that total-effect sensitivity indices, which account for total contribution of the inputs to variations in the model response, should be used to compare the significance of the model inputs in controlling the model output. *P*_*cp*_ has a larger *S*_*Ti*_ than 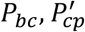, and 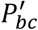, which shows *P*_*cp*_ is a more influential parameter in the model.

**Figure 5.**
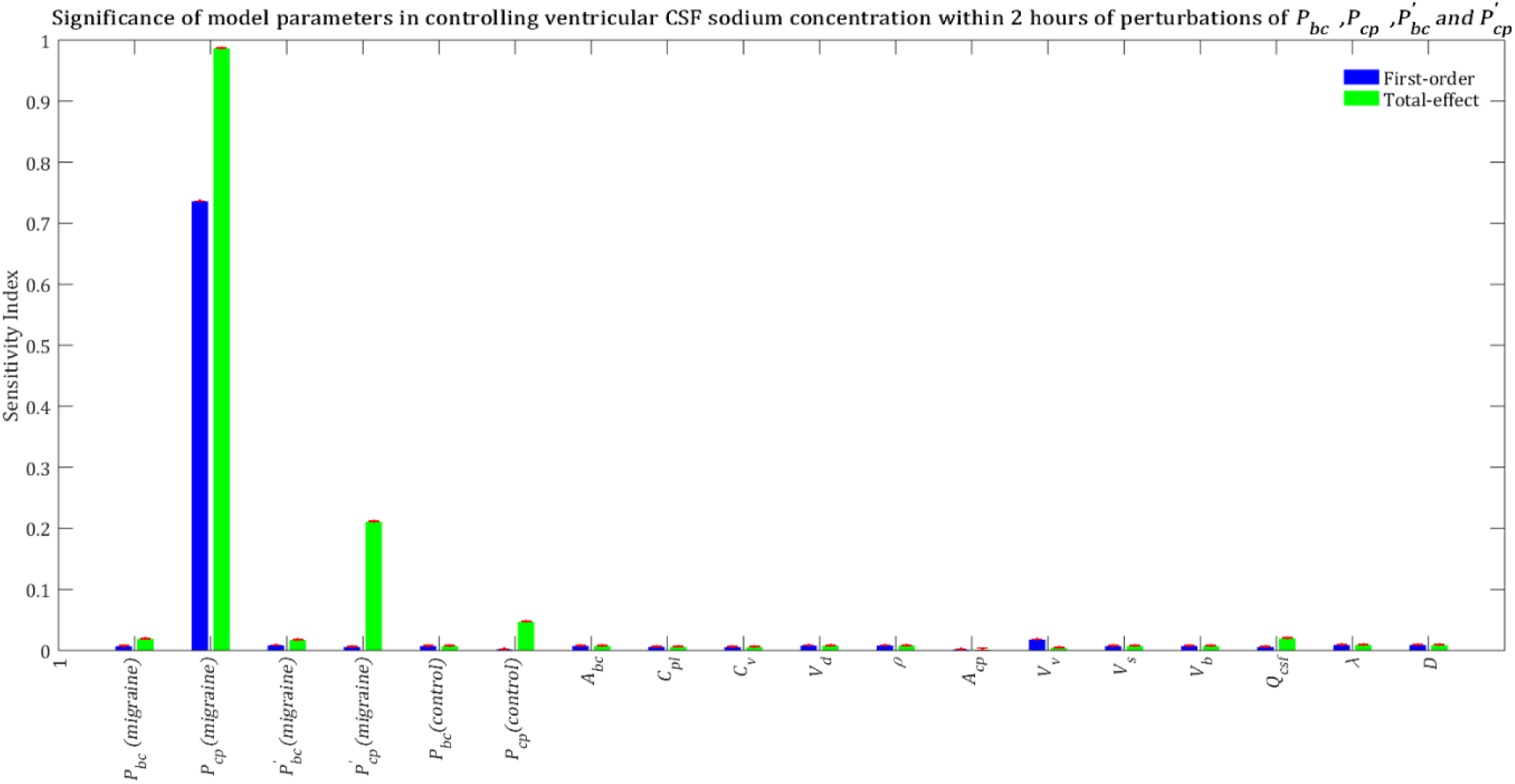
Sensitivity ranking of the model parameters. The model output was set to the time integral of *C*_*v*_ within 2 hours after perturbation of the model’s parameters. The blue bars represent first-order sensitivity indices, while the green bars show the total-effect sensitivity indices. The error bars, shown in red, indicate the bootstrap confidence intervals (95% confidence intervals) of the mean values.

Total-effect sensitivity indices of some of the parameters are smaller than 0.01 (Fig. 5). This means that the variations of these parameters do not influence the variance of the model output significantly; thus these parameters can be fixed at arbitrary values within their ranges [32, 33]. Figure 6 demonstrates the rank order of the model parameters when the model output was defined as the percent change of total subarachnoid sodium concentration within 2 hours after perturbations of control 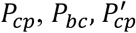, and 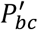 due to migraine triggers:

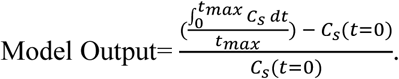

**Figure 6.**
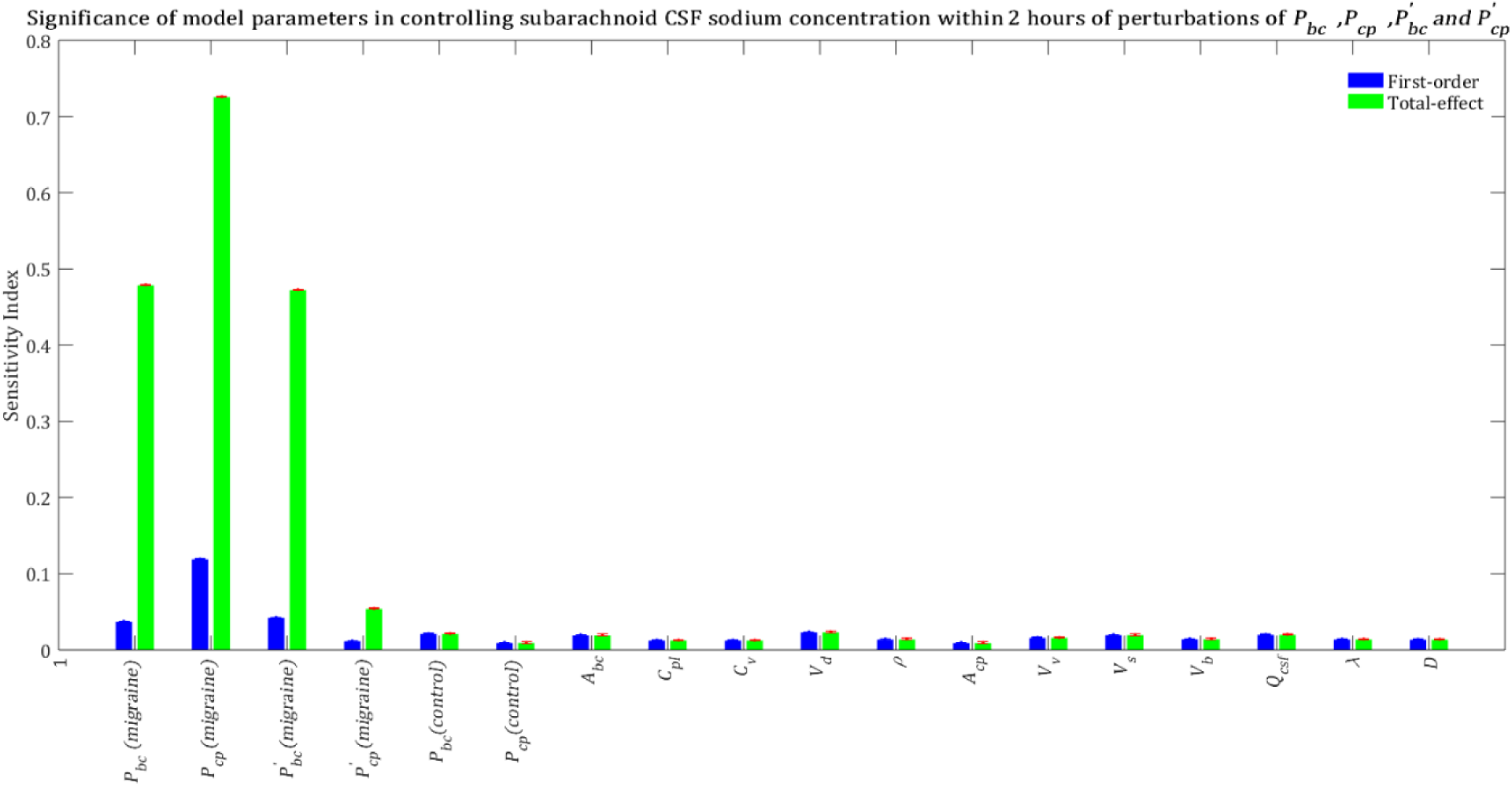
Relative significance of the model parameters in controlling subarachnoid CSF sodium concentration (*C*_*s*_) within 2 hours of the perturbation onset (*t*_*max*_ = 2 h). The blue bars represent first-order sensitivity indices, while the green bars show the total-effect sensitivity indices. The error bars, shown in red, indicate the bootstrap confidence intervals (95% confidence intervals) of the mean values.

Our results indicate that subarachnoid CSF sodium concentration is highly sensitive to pathophysiological changes in *P*_*cp*_, *P*_*bc*_ and 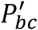 (Fig.6). The fact that pathophysiological variations of *P*_*bc*_ and 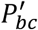 are more important in influencing subarachnoid sodium concentration than ventricular sodium concentration (Figs. 5-6) is because variations in *P*_*bc*_ and 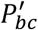 not only can affect sodium exchange at the contact surface of subarachnoid CSF and brain tissue, but also can influence sodium exchange between the ventricular system and brain tissue, thus affecting the amount of sodium entering the subarachnoid space from the ventricular system.

We also performed a GSA to identify the influential parameters when the model output was the percent change in total level of brain sodium after 2 hours of perturbations of the physiological 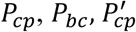, and 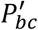 due to migraine triggers:

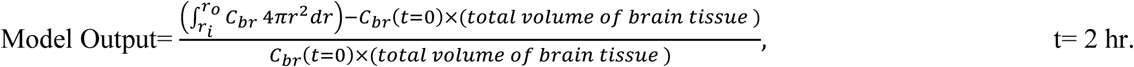

Our results demonstrate that brain tissue sodium level is highly sensitive to pathophysiological variations in 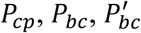 and 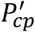 in order of decreasing sensitivity (Fig. 7). This result implies that sodium exchange between CSF and brain tissue at the contact surface of the ventricular system and brain tissue, as well as at the contact surface of the subarachnoid space and brain tissue can significantly influence brain sodium levels during migraine.

**Figure 7.**
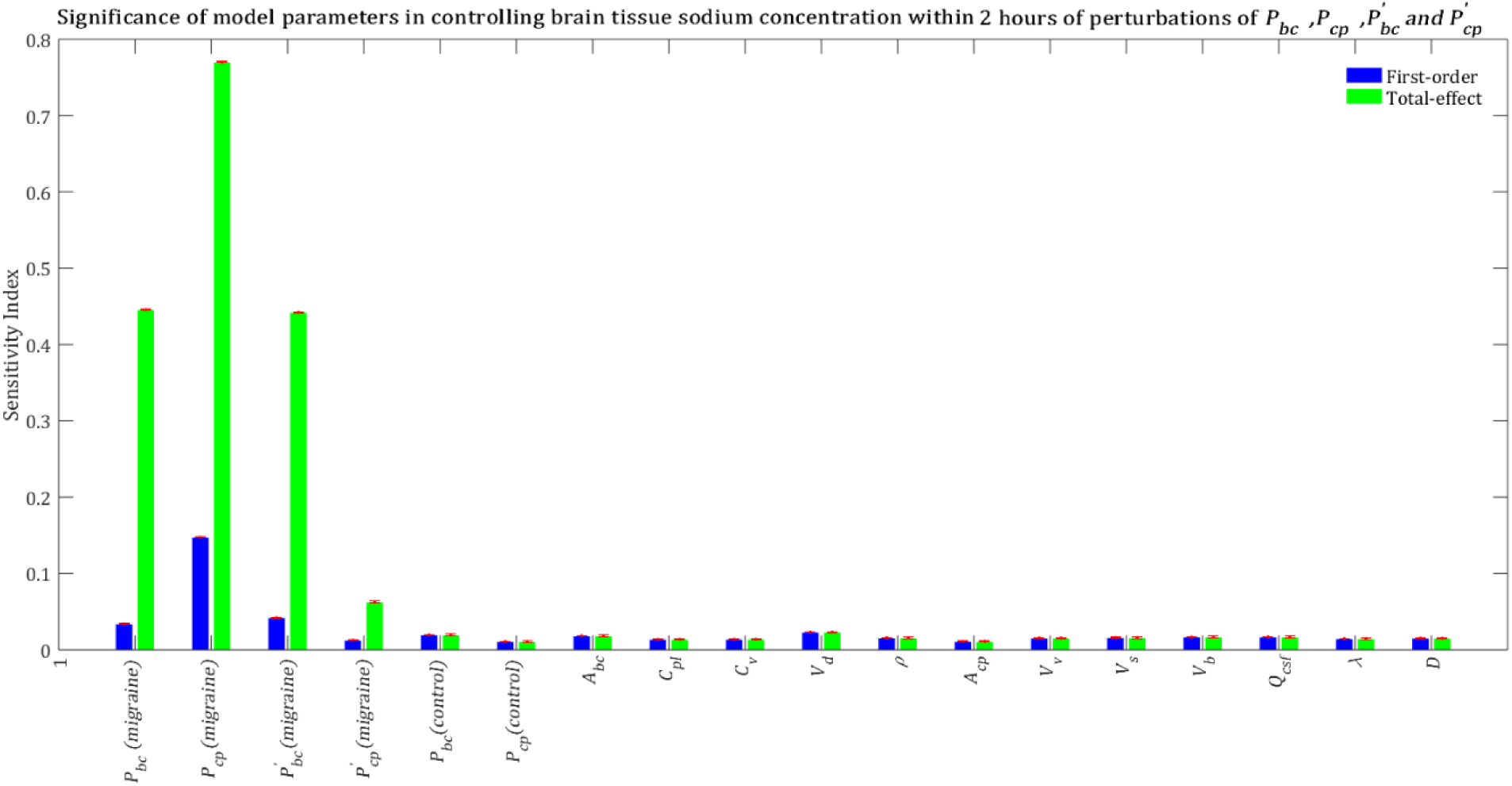
Relative importance of the model parameters in controlling brain tissue sodium levels within 2 hours of the perturbation onset (*t*_*max*_ = 2 h). The blue bars represent first-order sensitivity indices, while the green bars show the total-effect sensitivity indices. The error bars, shown in red, indicate the bootstrap confidence intervals (95% confidence intervals) of the mean values.

The above results were obtained after perturbing 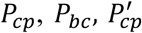, and 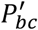 at t=0 and keeping them unchanged during the experiment time, i.e. t= 2 h. In order to investigate the impact of total experiment time on our results, we repeated our numerical experiments using different total experiment times including t= 10 min, t= 30 min and t= 1 h. Our results show that brain tissue, ventricular CSF and subarachnoid CSF sodium levels are mainly sensitive to pathophysiological variations in 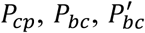 and 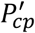 (See Supplementary Information: Figures S1-S9). The significance of pathophysiological changes of 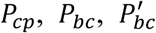 and 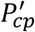 in influencing the ventricular CSF, subarachnoid CSF and brain tissue sodium levels at different total experiment times is shown in Table 2.

**Table 2.**
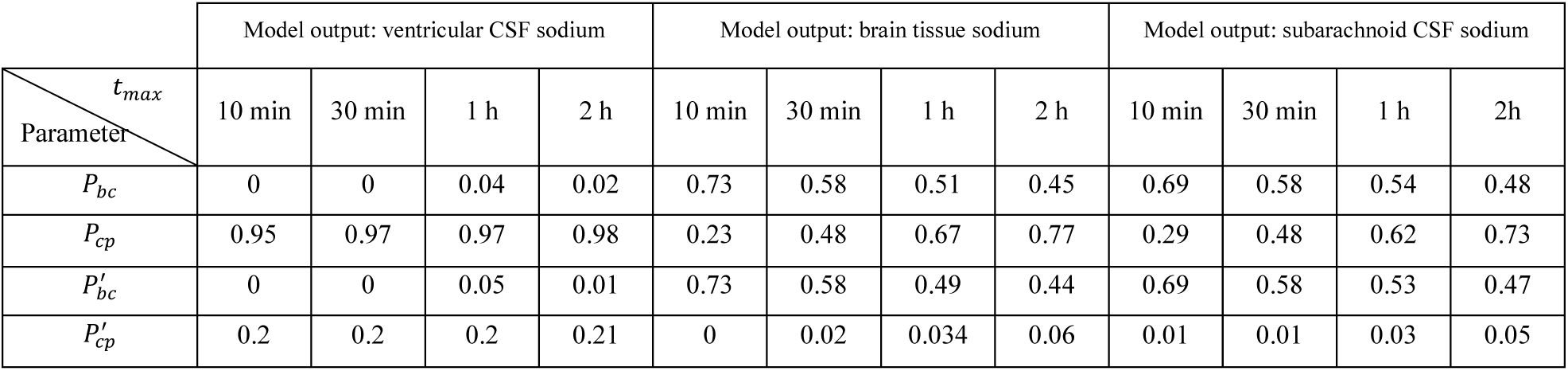
Total-effect sensitivity indices of the permeability coefficients at different total experiment times

| $t_{max}$<br>Parameter | Model output: ventricular CSF sodium | | | | Model output: brain tissue sodium | | | | Model output: subarachnoid CSF sodium | | | |
| --- | --- | --- | --- | --- | --- | --- | --- | --- | --- | --- | --- | --- |
|  | 10 min | 30 min | 1 h | 2 h | 10 min | 30 min | 1 h | 2 h | 10 min | 30 min | 1 h | 2h |
| $P_{bc}$ | 0 | 0 | 0.04 | 0.02 | 0.73 | 0.58 | 0.51 | 0.45 | 0.69 | 0.58 | 0.54 | 0.48 |
| $P_{cp}$ | 0.95 | 0.97 | 0.97 | 0.98 | 0.23 | 0.48 | 0.67 | 0.77 | 0.29 | 0.48 | 0.62 | 0.73 |
| $P'_{bc}$ | 0 | 0 | 0.05 | 0.01 | 0.73 | 0.58 | 0.49 | 0.44 | 0.69 | 0.58 | 0.53 | 0.47 |
| $P'_{cp}$ | 0.2 | 0.2 | 0.2 | 0.21 | 0 | 0.02 | 0.034 | 0.06 | 0.01 | 0.01 | 0.03 | 0.05 |

Our results demonstrate that the ventricular CSF sodium concentration is highly sensitive to pathophysiological variations in *P*_*cp*_, independent of experiment duration time. However, brain tissue and subarachnoid CSF sodium levels are more sensitive to pathophysiological variations of *P*_*bc*_ and 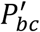 than that of *P*_*cp*_ at short total experiment times (such as 10 minutes and 30 minutes). Pathophysiological variations of *P*_*cp*_ become more important than variations of *P*_*bc*_ and 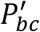 in controlling brain tissue and subarachnoid CSF sodium concentrations at longer experiment times (such as 1 hour and 2 hours). This implies that the BCSFB becomes more important in controlling brain tissue sodium homeostasis as time passes. This is because the ventricular CSF, whose sodium content is largely regulated by the BCSFB, would have enough time to influence sodium levels of its downstream compartments, including the brain tissue and the subarachnoid space.

To investigate the dynamics of sodium exchange between the CSF and brain tissue at the interface of brain tissue and the ventricular system, and at the contact surface of brain tissue and subarachnoid space during an episode of migraine, we randomly sampled 10^5^ sets of parameters, following a uniform distribution over a 18-dimensional parameter space and compared the average absolute sodium flux (*q*_*v*_) between brain tissue and ventricular CSF, with the average absolute sodium flux (*q*_*s*_) between the brain tissue and subarachnoid CSF. The average absolute fluxes *q*_*v*_ and *q*_*s*_ are defined by

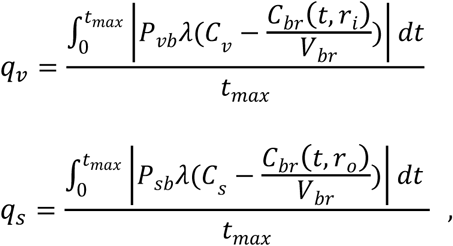

where *t*_*max*_ = 2 h. Figure 8 shows the ratio of *q*_*v*_ to *q*_*s*_ for the 10^5^ randomly sampled parameters. Our results indicate that the ratio of *q*_*v*_ to *q*_*s*_ is greater than 1 for the majority of the samples, which indicates that the absolute sodium flux at the interface of the ventricular system and the brain tissue is greater than the absolute sodium flux at the contact surface of the subarachnoid space and the brain tissue. Similar results were obtained for other total experiment times including *t*_*max*_ = 10 min, 30 min, 1 h (data not shown).

**Figure 8.**
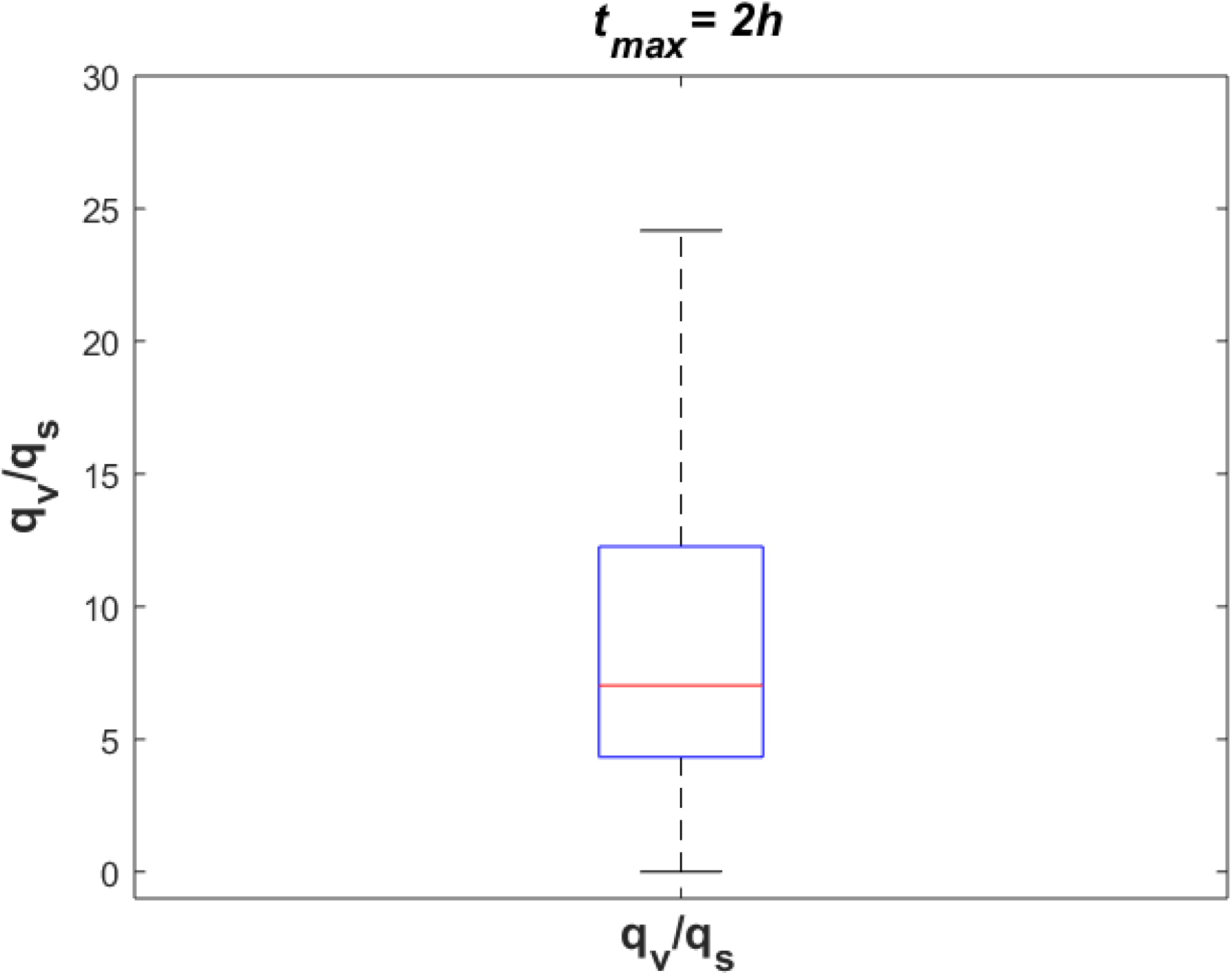
The ratio of absolute sodium flux at the interface of the ventricular system and the brain tissue (*q*_*v*_) to absolute sodium flux at the interface of the subarachnoid space and the brain tissue (*q*_*s*_). 10^5^ points were sampled randomly following a uniform distribution to generate this Figure.

## 4. Discussion

Previous studies [2, 3] have indicated that migraine sufferers have higher levels of CSF and brain tissue sodium than the control group. However, blood levels of sodium remain unchanged during migraine [2]. Under the hypothesis that these elevated sodium levels are due to variations in the influx and/or efflux permeability of the BCSFB and/or the BBB to sodium, we investigated the significance of variations in the influx and efflux permeabilities of the BCSFB and the BBB to sodium in influencing CSF and brain tissue sodium levels. In this regard, first we developed a computational model for sodium exchange between different brain compartments, i.e. blood, brain tissue, ventricular and subarachnoid CSF. The model presented in this paper is similar in some respects to that of Smith and Rapoport [8]. However, there are two major differences between our model and theirs. First, our model includes the ventricular system and subarachnoid space as separate compartments. Thus, our model can distinguish between the ventricular and subarachnoid CSF, as well as provide insight into the dynamics of sodium exchange between the CSF and brain tissue at the interface of brain tissue and the ventricular system, and at the contact surface of brain tissue and the subarachnoid space. Second, we have proposed a more realistic model of brain tissue compared to previous studies [8, 11, 35]. Unlike previous studies that modeled brain tissue as a rectangular sheet bathed on two opposite sides by CSF, we modeled brain tissue as the area between two concentric spheres. Concentric spheres are more similar to the real shape of a rat brain, which resembles an ellipsoid. As a result, the contact surface area of the brain tissue and the subarachnoid space is larger than that of the brain tissue and the ventricular system in our model. Thus, sodium exchange between the CSF and brain tissue at the two contact surfaces, as well as sodium diffusion in the brain tissue were modeled more accurately in this paper than in previous studies.

We performed a global sensitivity analysis to compare the significance of the BCSFB and the BBB in controlling CSF and brain sodium levels. Our results indicate that pathophysiological variations of the BCSFB influx permeability coefficient to sodium (*P*_*cp*_) are more important than variations of the BCSFB efflux permeability coefficient 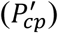, the BBB influx permeability coefficient (*P*_*bc*_) and the BBB efflux permeability coefficient 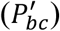 to sodium in controlling ventricular CSF sodium concentrations. Brain tissue and subarachnoid CSF sodium levels are more sensitive to pathophysiological variations of *P*_*bc*_ and 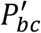 than to variations of *P*_*cp*_ when total experiment time is 10 minutes or 30 minutes, while *P*_*cp*_ becomes more important that *P*_*bc*_ and 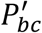 in influencing brain tissue and subarachnoid CSF sodium levels when total experiment time is 1 h or 2 h. Overall, our results show that *P*_*cp*_ plays an important role in the regulation of brain sodium homeostasis. *P*_*cp*_ represents the net movement of sodium from blood to CSF, which is regulated by a variety of BCSFB sodium transporters such as Na^+^, K^+^-ATPase [36-39], ENaC [36, 40] and NKCC1 [41]. Thus, variations in *P*_*cp*_ can be attributed to hyperactivity and/or hypoactivity of one or more of these sodium transporters. Our theoretical mechanism implies that the disturbed sodium homeostasis in the brain during a migraine is most likely due to overactivity of Na^+^, K^+^-ATPases at the BCSFB and the BBB [42]. This is in part because many regulators of Na^+^, K^+^-ATPase such as insulin [43, 44], dopamine [45, 46], glutamate [47], etc are involved in the pathophysiology of migraine (See [42] for a comprehensive review). Furthermore, there are several lines of evidence supporting that CSF secretion as well as sodium transport from the BCSFB cells, a.k.a choroid plexus epithelial cells, to CSF is mostly mediated by Na^+^, K^+^-ATPases, which are expressed on the apical membrane of the BCSFB cells [38, 48, 49]. It has been shown that intracerebroventricular infusion of ouabain, an Na^+^, K^+^-ATPase inhibitor, at 10 µg/day decreases CSF sodium concentration by almost 8 mM in Wister rats on a high-salt diet [37]. Ouabain can also reduce sodium transport from blood to CSF by 34% and 60% in frogs and rabbits, respectively [38, 39]. Thus, not only can the altered homeostasis of BCSFB Na^+^, K^+^-ATPases be a potential cause of the elevated CSF sodium concentration in a migraine, but also BCSFB Na^+^, K^+^-ATPase could be a candidate drug target to correct the elevated levels of sodium in CSF of migraine sufferers, potentially treating migraine. This prediction needs to be tested experimentally for different migraine triggers. ENaC is another sodium transporter which can play a key role in the regulation of CSF sodium levels. Overexpression of ENaC at the basolateral membrane of the BCSFB cells may increase *P*_*cp*_. It has been shown that overexpression of ENaC at the basolateral membrane of BCSFB increases sodium transport from blood to BCSFB cells in salt-induced hypertension in a mouse model of Liddle’s Syndrome [40]. Sodium transported into BCSFB cells is pumped out into CSF by Na^+^, K^+^-ATPases, leading to an increase in CSF sodium concentration. In Wister rats, ENaC is expressed at both membranes of BCSFB cells with a higher density at the CSF-facing (apical) membrane compared to the basolateral membrane [50, 51]. This suggests that ENaC may play a major role in sodium uptake from CSF into BCSFB cells [52]. It should be noted that sodium movement through ENaC is likely to be unidirectional; thus, variations in the activity levels of ENaC at the apical membrane of BCSFB cells can potentially change 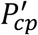, while variations in ENaC activity levels at the basolateral membrane of BCSFB cells can possibly change *P*_*cp*_. It is not known how the expression levels of ENaC on the different membranes of BCSFB cells are affected by migraine triggers. The other main BCSFB sodium transporter is NKCC1, which can regulate CSF production [53] and sodium movement from blood to CSF [41]. Overall, sodium transport from blood to CSF across the BCSFB is regulated by a variety of transporters, channels and proteins, whose interactions with each other are not well understood. Further experimental studies are needed to elucidate the potential effects of various migraine triggers on the activity and expression levels of BCSFB Na^+^, K^+^-ATPase, ENaC and NKCC1.

Our results suggest that the BBB can play a more important role than the BCSFB in the regulation of brain tissue and subarachnoid CSF sodium concentrations within 30 minutes of pathophysiological perturbations of 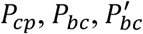 and 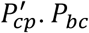 and 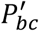 were used in the current model to simulate the net movement of sodium from blood to brain tissue, and from brain tissue to blood, respectively. Variations in *P*_*bc*_ and 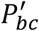 can be attributed to altered homeostasis of the transporters, which mediate sodium movement across the BBB. The principal routes for sodium entry across the luminal membrane of the BBB endothelial cells are likely to be NKCC1 [54, 55] and NHE1,2 [56], while sodium is mainly pumped out of the BBB endothelial cells into brain ISF by Na^+^, K^+^-ATPase [7, 57, 58]. It has been suggested that sodium transport from the brain ISF into the BBB endothelial cells is mainly mediated by sodium-linked transporters of organic solutes, including those for amino acids [7]. NHE 1,2 can also potentially contribute to sodium entry across the abluminal membrane of endothelial cells. However, the impacts of migraine triggers on the activity and expression levels of these sodium transporters are yet to be understood. Our results suggest that alterations of BBB sodium transporters homeostasis have more significant effects than variations of BCSFB sodium transporters homeostasis on brain tissue sodium levels within 30 minutes of the perturbation onset. It should be noted that our results were obtained using GSA, which gives us some insight into the importance of influx and efflux permeability of the BCSFB and the BBB to sodium in controlling CSF and brain tissue sodium by covering the entire parameter space, where all model parameters can vary within the specified ranges. Thus, in a rat model, the intrinsic variations between a population of rats of the same type were considered in this work.

This study has some limitations. First, for simplicity, we modeled the rat brain with three spheres. However, the real geometry of a rat brain is more complicated. A more realistic model of the brain and ventricles can provide a better understanding of the phenomenon under study. Second, we modeled the CSF with two well-mixed compartments, i.e. the ventricular system and the subarachnoid space. However, CSF flows through the lateral ventricles, the third ventricle, the cerebral aqueduct, the fourth ventricle, the cisterns and the subarachnoid space. Sodium concentration can vary slightly to significantly from one ventricle to another one and to the subarachnoid space. Thus, the current model can be improved to include all of the ventricles and subarachnoid space as separate compartments. To do this, further information regarding the dynamics of sodium transport between different ventricles and adjacent brain tissues is needed. Furthermore, we assumed that there is no rate-limiting diffusion between the CSF and brain tissue at the two contact surfaces of the CSF and brain tissue. This results in instantaneous equilibrium between CSF sodium concentration and brain ISF sodium concentration at the contact surface of brain tissue and CSF [8]. This assumption may not be true for some ependymal regions such as those in the third ventricle as it has been shown that benzamil, an ENaC blocker can prevent sodium movement from the third ventricle CSF into brain tissue across the ependyma [52]. Third, for simplicity we assumed that the value of the sodium distribution factor (*V*_*d*_) remains constant after perturbations of the BCSFB and the BBB permeability coefficients to sodium. Thus, we estimated the ISF sodium concentration by 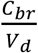 (Eq. 4). This assumption implies that the ratio of extracellular sodium concentration to intracellular sodium concentration remains unchanged at any time after perturbations of the permeability coefficients. In other words, sodium is always distributed between the ISF and the brain cells in the ratio of their physiological sodium contents. Previous studies made a somewhat similar assumption to estimate the ISF sodium concentration from brain tissue sodium levels, using the cerebral distribution volume of sodium [8]. The physiological value of *V*_*d*_ was found to be 0.34 ml/g using the average physiological ISF sodium concentration of 145 mM [14] and the average brain tissue sodium concentration of 50 mM (=50 × 10^−6^ mol/g) [59]. The obtained value of 0.34 ml/g for *V*_*d*_ in this work is the same as the value of the cerebral distribution volume of sodium [8]. The dynamics of sodium exchange between the brain cells and the ISF can be better understood by adding the brain cells as a new compartment to the current model. Our model can be expanded to include brain cells once more information becomes available regarding the permeability coefficients of brain cells to sodium. Fourth, we perturbed 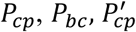, and 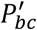 at t=0 and kept them unchanged during the experiment time. However, in reality the BCSFB and the BBB permeability coefficients likely change over time. Thus, the model presented in this study can be used to study the contribution of the BCSFB and the BBB to variations in the brain tissue and CSF sodium concentrations once there is more information about time-dependent variations of the BBB and BCSFB permeabilities to sodium during an episode of migraine with a particular trigger. Fifth, we assumed that diffusion is the major mechanism of sodium movement in the brain tissue. Although there are several lines of evidence supporting the existence of a convective transport mechanism called the glymphatic system in the brain [12, 13], several aspects of glymphatic circulation, including whether interstitial transport is propagated by convective flow or diffusion [60, 61], the identity of the ISF bulk flow driving forces [62, 63], and the role of astrocyte water permeability/aquaporin4 [61] are still controversial. Furthermore, it is not well understood how the proposed transport mechanisms are affected during migraine and how these mechanisms interact with the BBB to regulate ionic homeostasis in the brain. In this work, we ignored sodium transport between the CSF and brain ISF via convection, as it has been shown that diffusion (without convection) in the brain tissue is enough to account for many experimental transport studies in the brain parenchyma [61].

Intuitively, we think that adding the convective CSF transport from the subarachnoid space to the brain ISF, based on the proposed glymphatic circulation, increases the effects of subarachnoid CSF (in general CSF) on the brain tissue sodium levels, as the convective transport mechanism allows more sodium to be transported in a shorter amount of time compared to diffusive transport. Thus, the BCSFB would become more important in controlling brain tissue sodium levels. However, the exact extent of the contribution of the glymphatic system to the regulation of brain sodium homeostasis depends on not only the dynamical properties of the glymphatic system such as the rate of glymphatic flow, the glymphatic efflux pathways and the ISF bulk flow driving force, but also the dynamic interactions between the glymphatic flow, the BBB and brain diffusive transport mechanisms. The current model can be expanded to include the convective CSF flow from the subarachnoid space to brain ISF once more information regarding the contribution of the glymphatic flow to the regulation of brain sodium homeostasis becomes available. Finally, we ignored water fluxes between the model compartments. Thus, the volumes of the model compartments remain unchanged during the experiment time. This is because variations of the permeability coefficients within the specified ranges in this study result in gradual changes in the brain ISF sodium concentration, which suggests that the ISF osmolality changes gradually. The gradual variations in the ISF osmolality give the brain cells enough time to adjust to the changes in the extracellular space; so that they can minimize the variations in their volume through regulating the influx and efflux of osmotically active solutes between the intracellular and extracellular fluids. Previous in vitro studies showed that the cultured cerebellar neurons and C6 rat glioma cells can exhibit isovolumetric regulation when the extracellular osmolality changes at a rate less than or equal to 1.8 and 3 mOsmol/kg/min, respectively [64, 65]. The maximum possible rate of change of ISF sodium concentration in this work is 1.5 mM/min, equivalent to 1.5 mOsmol/kg/min. Thus, we believe that the brain cells, which make up 80% of total volume of the brain can significantly maintain their volume under the assumptions/conditions in our numerical simulation. This argument is in agreement with another experimental observation which suggests that a 50% decrease in the activity levels of Na^+^, K^+^-ATPase on the brain microvessels does not change total water content in the brain significantly [66]. Assuming that the brain tissue volume remains almost unchanged in this work, one can conclude that the CSF volume remains almost constant due to the rigid confines of the skull. We have also assumed that the CSF secretion rate remains unchanged after pathophysiological variations of the influx and/or efflux permeability coefficients of the BCSFB to sodium. Although it has been suggested that there is a positive correlation between the CSF secretion and sodium transport rates across the BCSFB [7], it is not known how and to what extent water movement is linked to sodium transport in the BCSFB during migraine. Migraine is accompanied with a complex chain of biochemical changes in the CSF and brain which may contribute, together with sodium, to regulation of water movement across the BCSFB. For instance, it has been shown that CSF (and plasma) content of organic osmolytes such as taurine and glutamate, which can significantly regulate brain cell volume homeostasis [67, 68], changes during migraine [69-72]. However, it is yet to be determined how the variations in organic osmolyte levels can influence the osmotically driven water transport across the choroid plexus. Thus, future experimental studies are needed to explore whether/how/to what extent water movement rate depends on sodium transport rate during migraine. The results presented in this work may vary depending on how the CSF flow rate changes during migraine. The current model can be extended to include dynamic water movement across the BCSFB once further information regarding the extent to which water movement is linked to sodium transport during migraine becomes available

## Conclusions

Our proposed mechanism for migraine suggests that a disturbance in brain sodium homeostasis causes migraine [42]. This sodium dysregulation is most likely due to variations in the influx and/or efflux permeability of the BCSFB and/or the BBB to sodium. The influx and efflux permeability of the BCSFB and the BBB to sodium represent the net effect of all transporters, channels and enzymes which contribute to movement of sodium across the interfaces. Thus, variations of the permeability coefficients can be caused by altered homeostasis of one or some of the sodium transport mechanisms at the interfaces. Unfortunately, understanding migraine pathophysiology is difficult, not only because the effects of various triggers on permeability of the BCSFB and the BBB to sodium are not known, but also because migraines have different triggers in different people. To approach this problem, we used mechanistic modeling together with global sensitivity analysis (GSA) to assess the relative importance of the BCSFB and the BBB in controlling CSF and brain tissue sodium levels. GSA provides insight into the significance of the BCSFB and the BBB in the regulation of brain sodium concentration when the exact extents of variations in the influx and efflux permeability coefficients of the BCSFB and the BBB to sodium are unknown. Our results show that the ventricular CSF sodium concentration is highly influenced by pathophysiological variations in the influx permeability coefficient of the BCSFB to sodium. Brain tissue and subarachnoid CSF sodium levels are more sensitive to pathophysiological variations in the BBB permeability coefficients than the BCSFB permeability coefficients to sodium at shorter total experiment times (such as 10 and 30 minutes), while the BCSFB becomes more important that the BBB in influencing total brain tissue and subarachnoid CSF sodium levels at longer experiment times (such as 1 and 2 h). These results suggest that the efficacy of different migraine treatment strategies may depend on the time elapsed from migraine onset. This prediction needs to be tested experimentally for different models of migraines. This study prompts the hypothesis that increased influx permeability of the BCSFB to sodium caused by altered homeostasis of the enzymes which transport sodium from blood to CSF is the potential cause of elevated brain sodium levels in migraines. This hypothesis needs to be tested experimentally. The current model can be used to simulate sodium transport across the BBB, the BCSFB and the ependymal surfaces for a particular migraine trigger, given that the effects of the migraine trigger on the BBB and the BCSFB permeabilities are known. Further studies on the activity levels of different BCSFB and BBB sodium transporters during migraine episodes with different triggers can help better understand migraine pathophysiology.

## Supporting information

Animation 1

Animation 2

Animation 3

Animation 4

## Declarations

### Consent for publication

Not applicable.

### Availability of data and material

All data generated or analyzed during this study are included in this published article.

### Ethics approval and consent to participate

Not applicable.

### Authors’ contributions

HG conceived and designed the experiments. HG performed the experiments. HG, SCG, LRP and MGH analyzed the results. HG, SCG, LRP and MGH wrote and approved the paper.

## Acknowledgment

Not applicable.

## Abbreviations

CSF: Cerebrospinal fluid
BBB: Blood-brain barrier
BCSFB: Blood-CSF barrier
ISF: Interstitial fluid
GSA: Global sensitivity analysis

## Supplementary Information

### Global Sensitivity Analysis

In this work, we used a MATLAB toolbox called SAFE [25] to perform a global sensitivity analysis (GSA). SAFE implements several GSA methods such as the Elementary Effects Test, Regional Sensitivity Analysis, and Sobol’s technique. Sobol’s method is a variance-based global sensitivity analysis technique which evaluates the sensitivity of the solutions with respect to the model parameters as well as the interactions between different parameters. Using the principles of variance decomposition, Sobol’s method ranks the parameters in terms of their importance. Given an integrable function *f* over a *p*-dimensional parameter space Ω^*p*^,

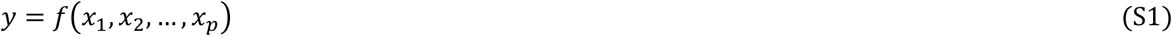

Each parameter can vary within a finite range. Sobol’s method considers expansion of the response into a set of functions of increasing dimensionality,

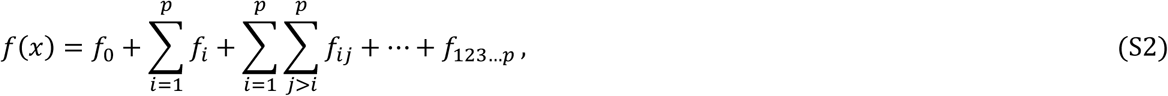

where each individual term is a function of the parameters in its index. The total variance of the function output is defined by

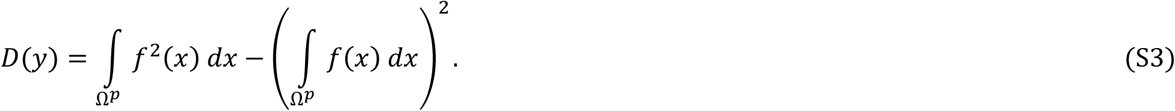

Sobol’s technique is based on decomposition of the total variance *D* into partial variances indicating the contributions from effects of individual parameters and combined effects of pairs of parameters. This decomposition is accomplished using the expansion of *f* into terms of increasing dimensions (Eq. S2),

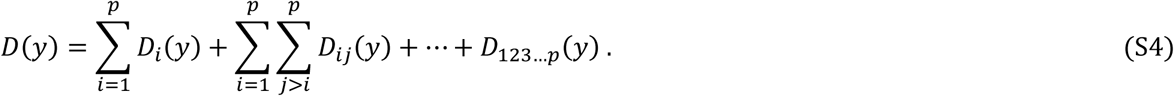

According to Sobol’s method, the first-order sensitivity index for each parameter is given by

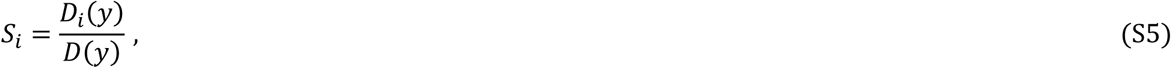

The first-order sensitivity index accounts for the main individual contribution of each model parameter to the variance of the model output. The Sobol’s total-effect index, on the other hand, represents total contribution of the input to the response variation. The total-effect index for parameter *x*_*i*_ is calculated by the sum of all sensitivity indices which have *i* in their index

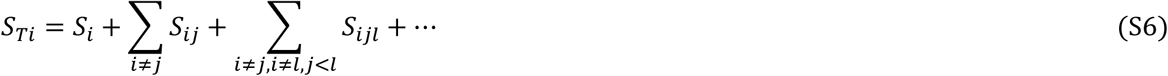

Based on Sobol’s approach, the necessary and sufficient condition for parameter *x*_*i*_ to be a noninfluential factor is *S*_*Ti*_ = 0. However, previous studies have indicated that a parameter can be considered noninfluential if its total-effect sensitivity index is smaller than 0.01, and significantly smaller than total-effect sensitivity indices of the rest of the parameters [32, 33, 73, 74].

---

**Figure S1.**
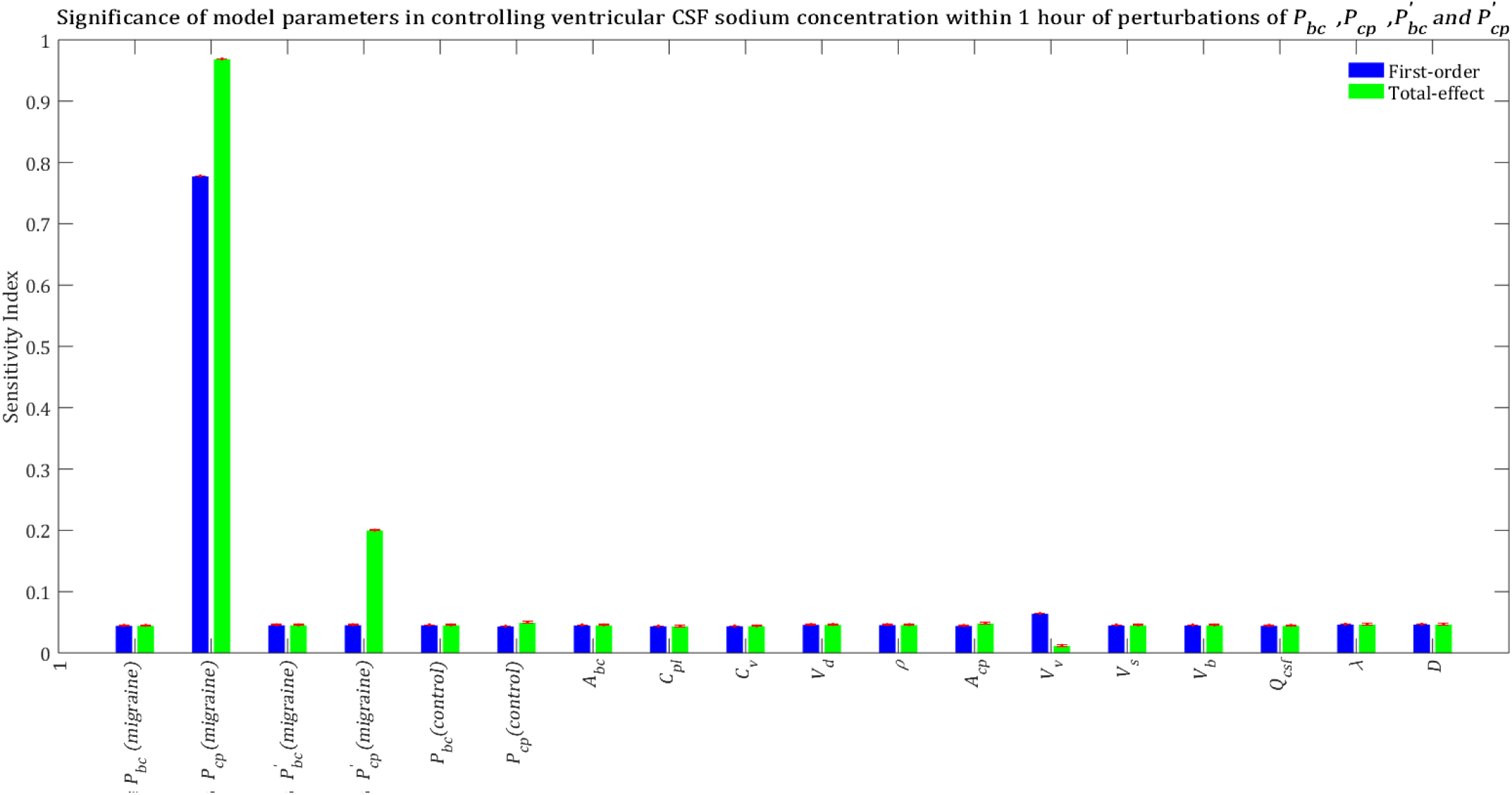
Relative significance of the model parameters in controlling ventricular CSF sodium concentration (*C*_*s*_) within 1 hour of the perturbation onset (*t*_*max*_ = 1 h). The blue bars represent first-order sensitivity indices, while the green bars show the total-effect sensitivity indices. The error bars, shown in red, indicate the bootstrap confidence intervals (95% confidence intervals) of the mean values.

**Figure S2.**
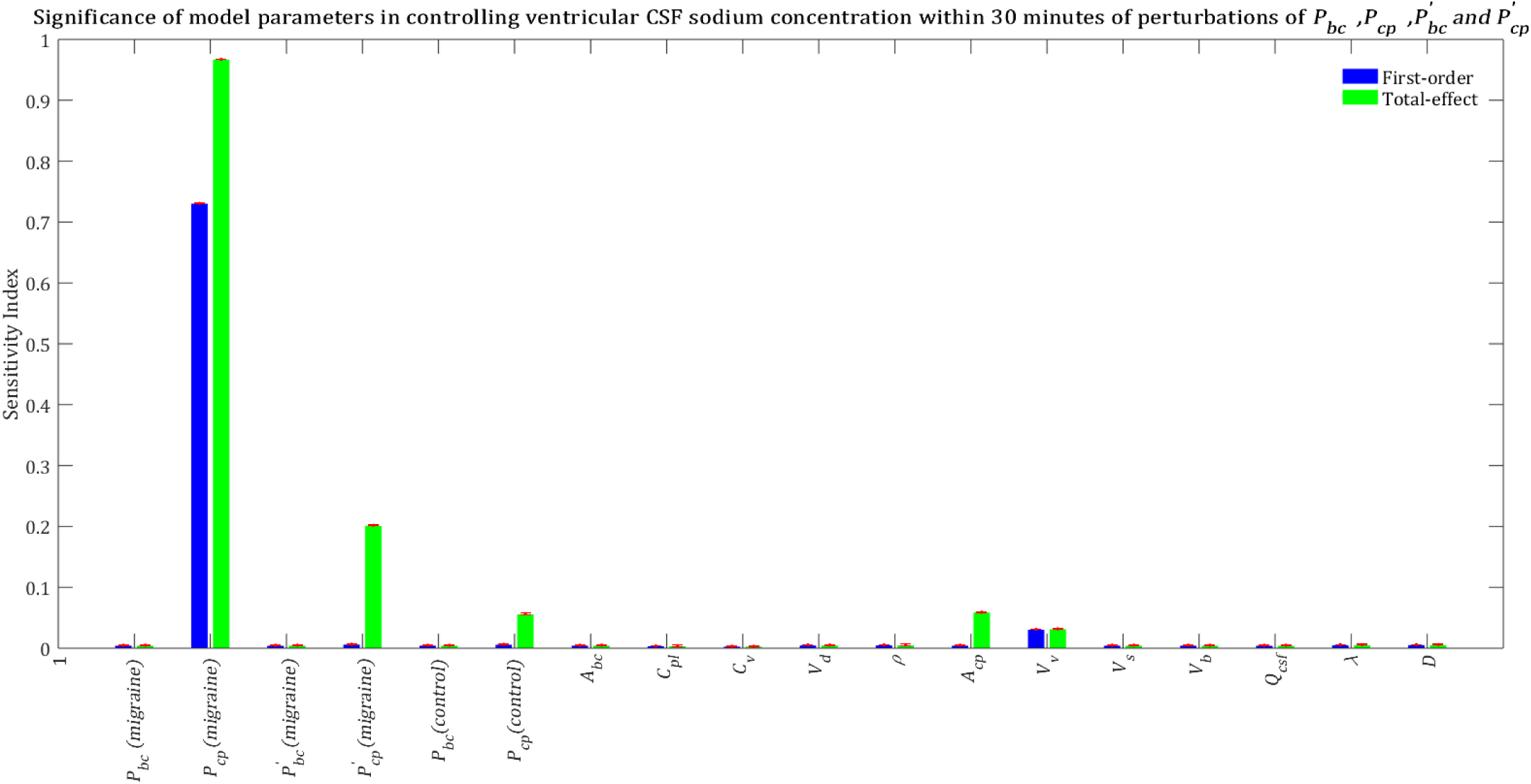
Relative significance of the model parameters in controlling ventricular CSF sodium concentration (*C*_*s*_) within 30 minutes of the perturbation onset (*t*_*max*_ = 30 m). The blue bars represent first-order sensitivity indices, while the green bars show the total-effect sensitivity indices. The error bars, shown in red, indicate the bootstrap confidence intervals (95% confidence intervals) of the mean values.

**Figure S3.**
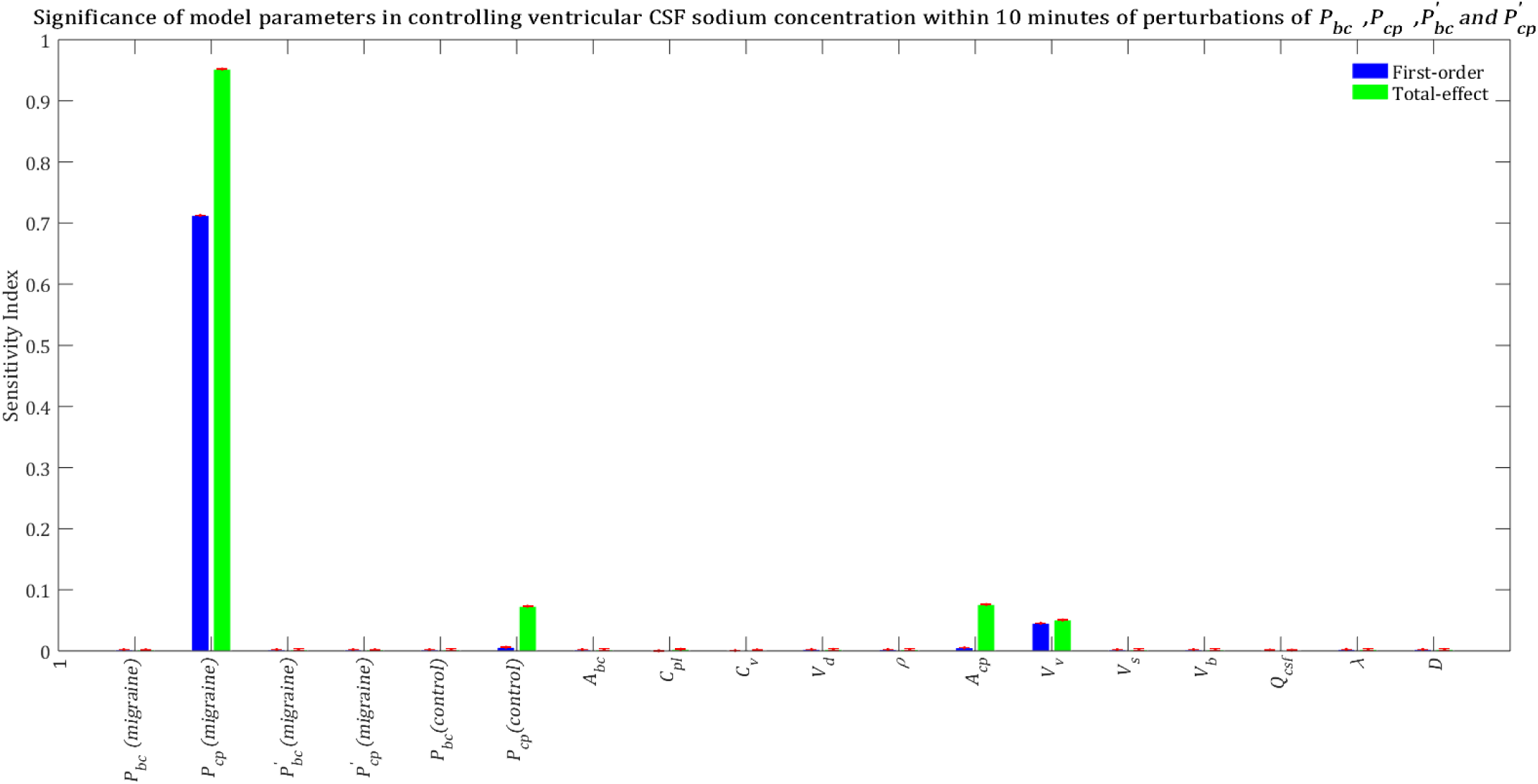
Relative significance of the model parameters in controlling ventricular CSF sodium concentration (*C*_*s*_) within 10 minutes of the perturbation onset (*t*_*max*_ = 10 min). The blue bars represent first-order sensitivity indices, while the green bars show the total-effect sensitivity indices. The error bars, shown in red, indicate the bootstrap confidence intervals (95% confidence intervals) of the mean values.

**Figure S4.**
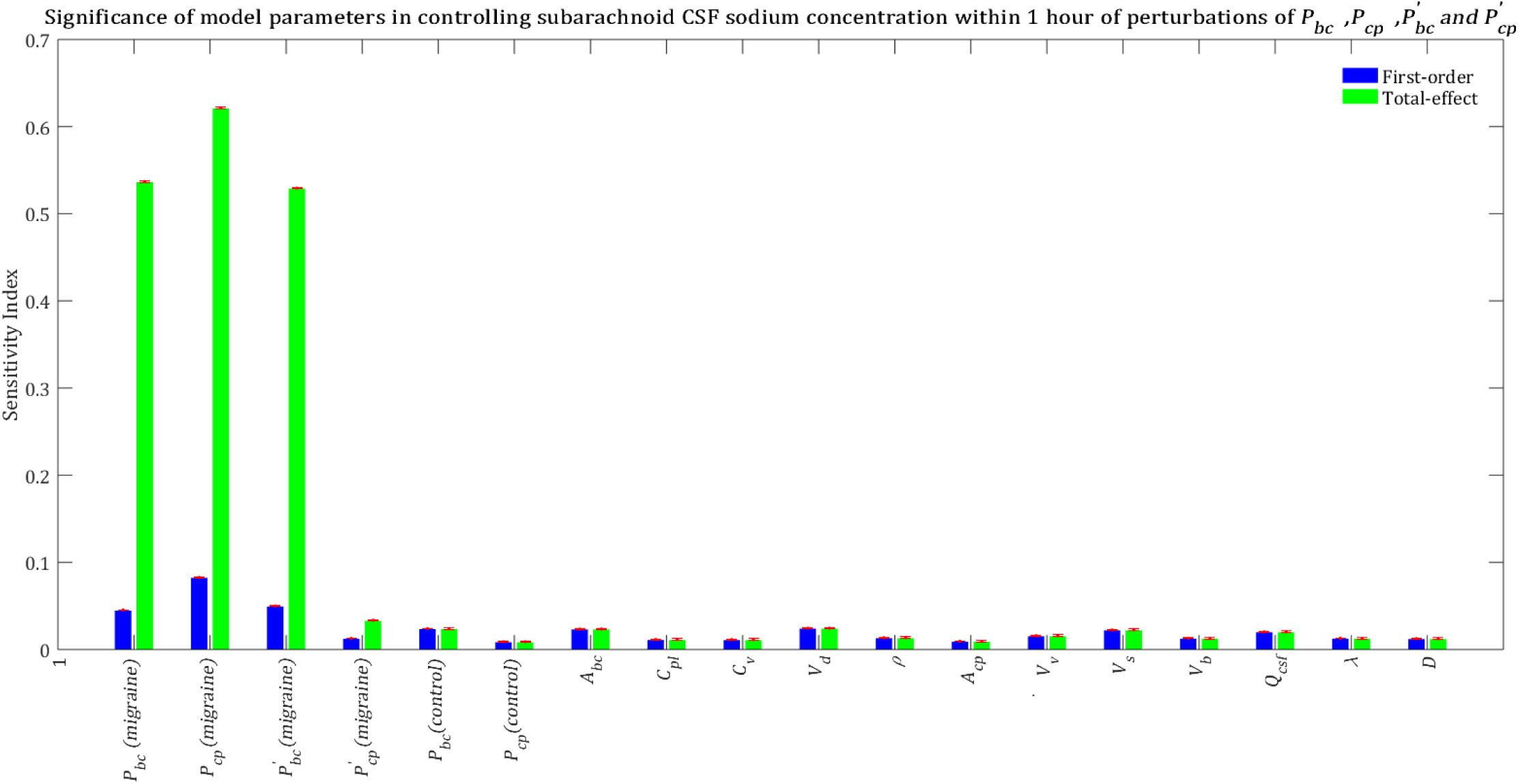
Relative significance of the model parameters in controlling subarachnoid CSF sodium concentration (*C*_*s*_) within 1 hours of the perturbation onset (*t*_*max*_ = 1 h). The blue bars represent first-order sensitivity indices, while the green bars show the total-effect sensitivity indices. The error bars, shown in red, indicate the bootstrap confidence intervals (95% confidence intervals) of the mean values.

**Figure S5.**
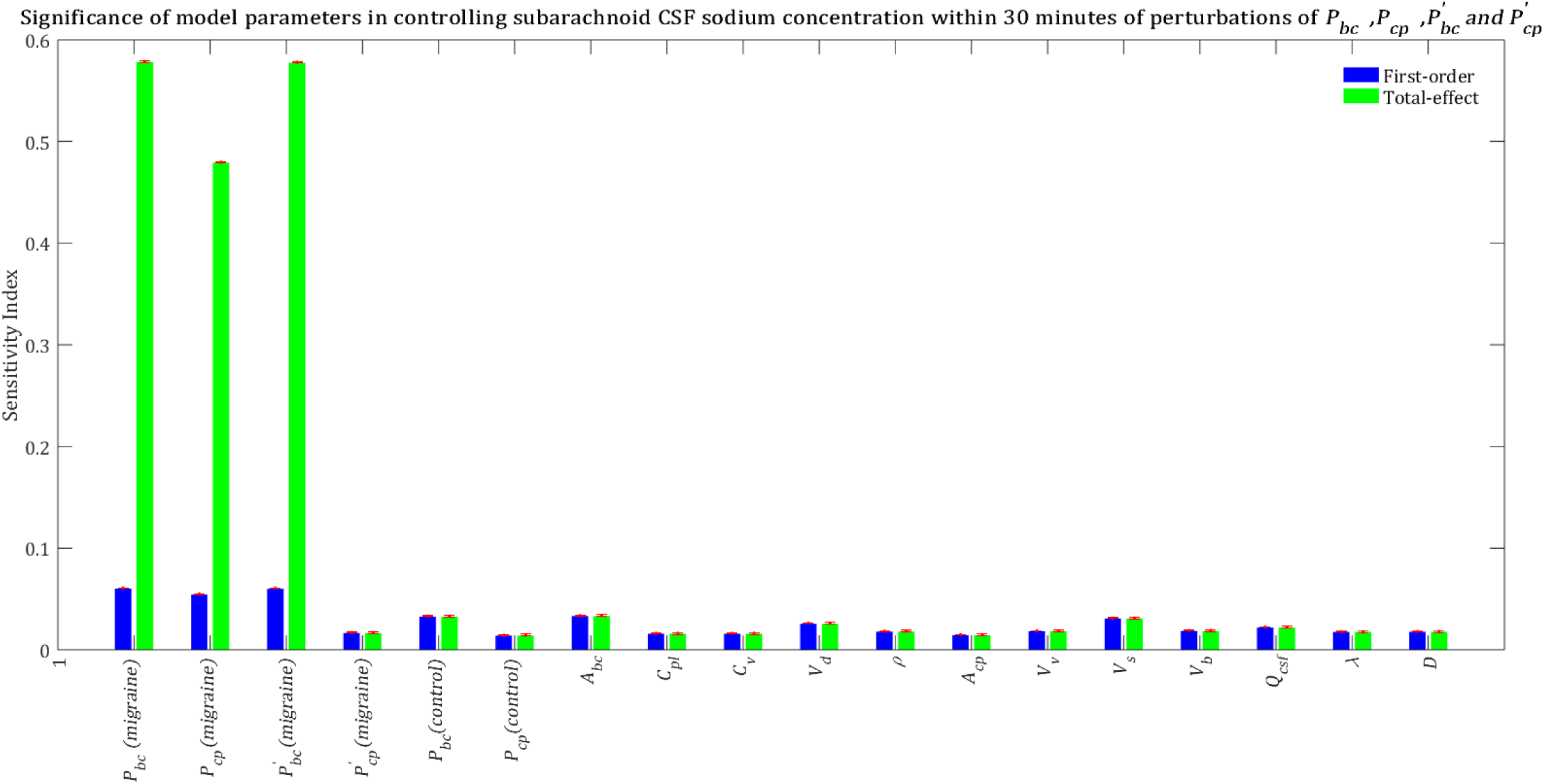
Relative significance of the model parameters in controlling subarachnoid CSF sodium concentration (*C*_*s*_) within 30 minutes of the perturbation onset (*t*_*max*_ = 30 min). The blue bars represent first-order sensitivity indices, while the green bars show the total-effect sensitivity indices. The error bars, shown in red, indicate the bootstrap confidence intervals (95% confidence intervals) of the mean values.

**Figure S6.**
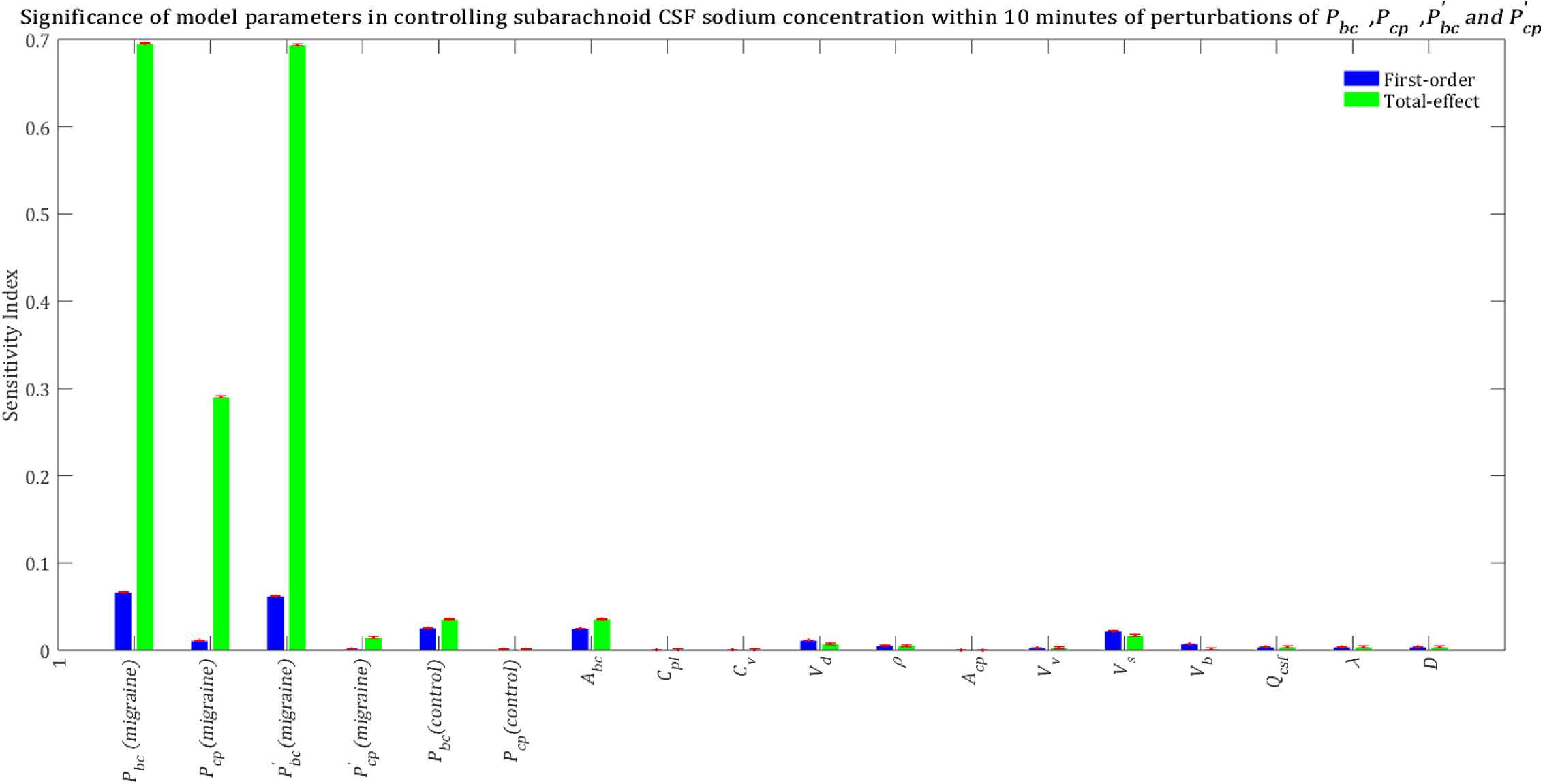
Relative significance of the model parameters in controlling subarachnoid CSF sodium concentration (*C*_*s*_) within 10 minutes of the perturbation onset (*t*_*max*_ = 10 min). The blue bars represent first-order sensitivity indices, while the green bars show the total-effect sensitivity indices. The error bars, shown in red, indicate the bootstrap confidence intervals (95% confidence intervals) of the mean values.

**Figure S7.**
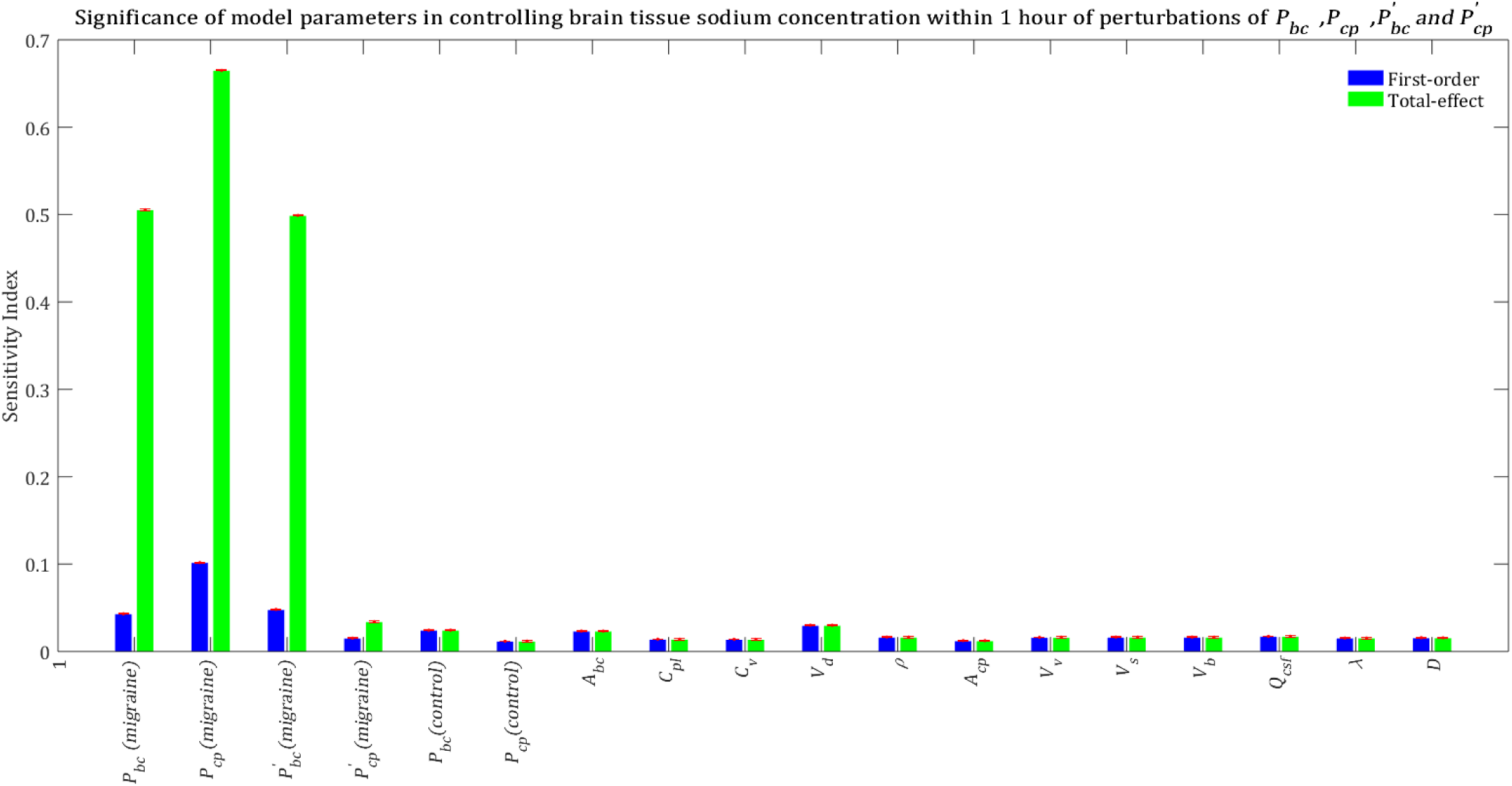
Relative importance of the model parameters in controlling brain tissue sodium levels within 1 hour of the perturbation onset (*t*_*max*_ = 1 h). The blue bars represent first-order sensitivity indices, while the green bars show the total-effect sensitivity indices. The error bars, shown in red, indicate the bootstrap confidence intervals (95% confidence intervals) of the mean values.

**Figure S8.**
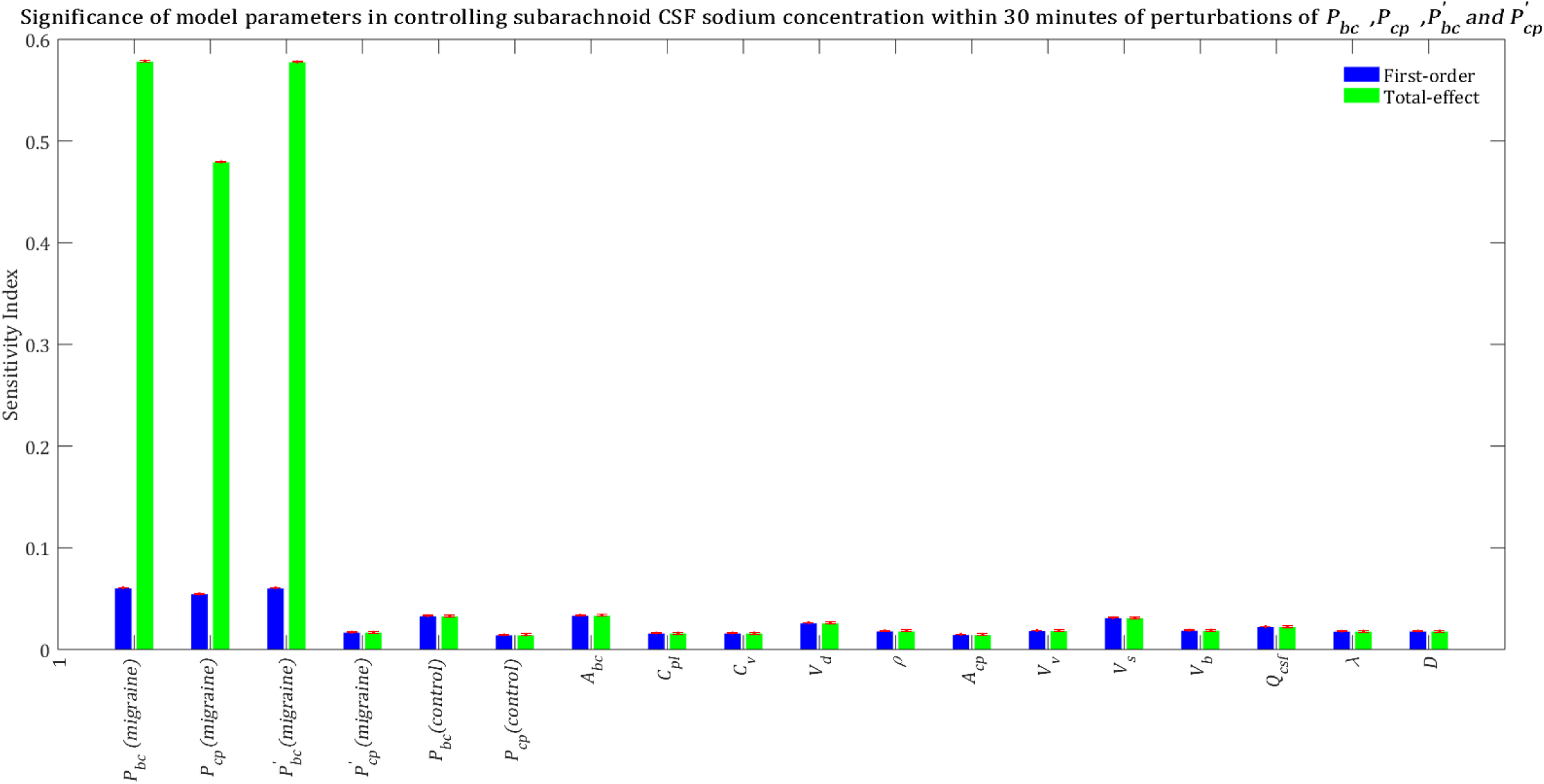
Relative importance of the model parameters in controlling brain tissue sodium levels within 30 minutes of the perturbation onset (*t*_*max*_ = 30 min). The blue bars represent first-order sensitivity indices, while the green bars show the total-effect sensitivity indices. The error bars, shown in red, indicate the bootstrap confidence intervals (95% confidence intervals) of the mean values.

**Figure S9.**
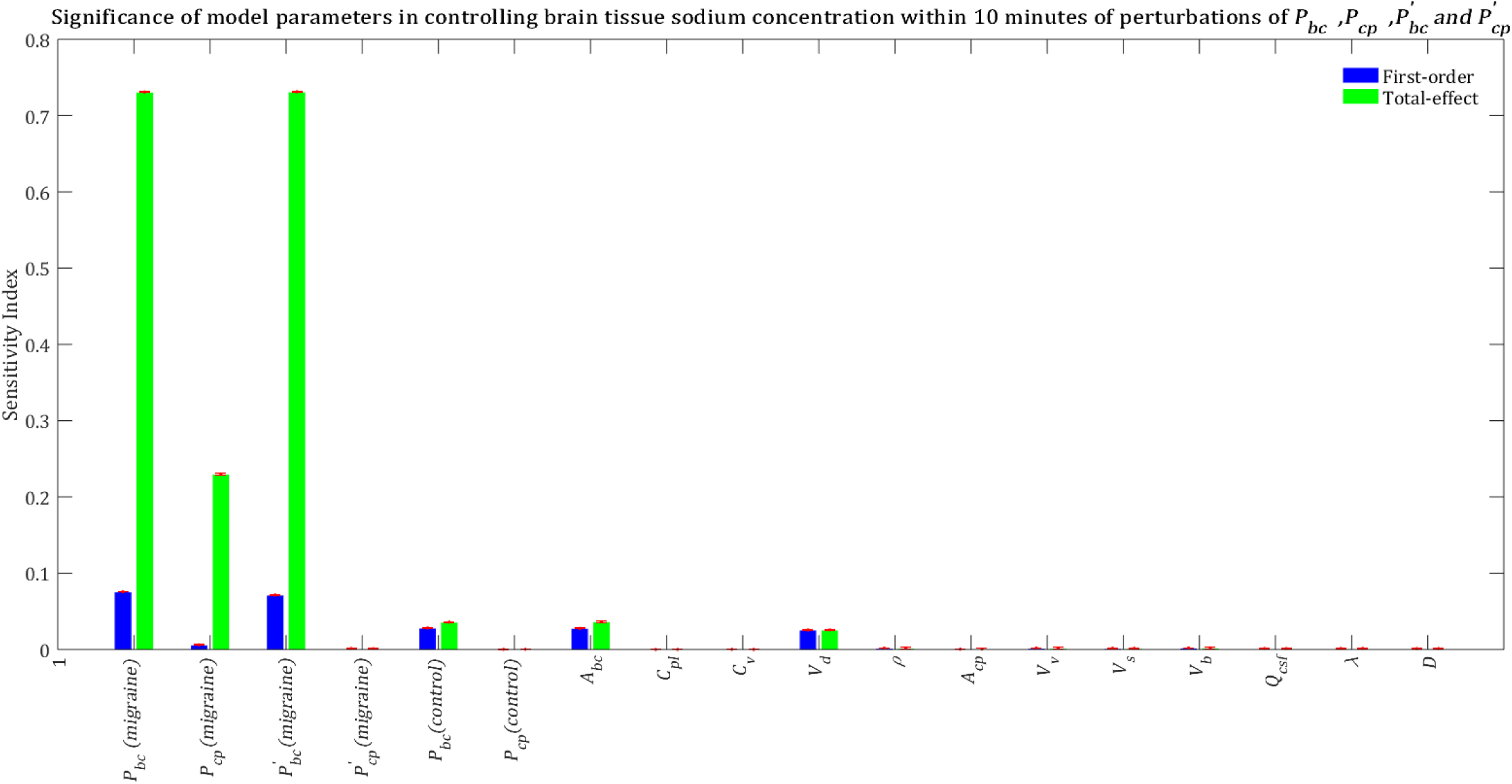
Relative importance of the model parameters in controlling brain tissue sodium levels within 10 minutes of the perturbation onset (*t*_*max*_ = 10 min). The blue bars represent first-order sensitivity indices, while the green bars show the total-effect sensitivity indices. The error bars, shown in red, indicate the bootstrap confidence intervals (95% confidence intervals) of the mean values.

